# Autonomous AI Agents Discover Aging Interventions from Millions of Molecular Profiles

**DOI:** 10.1101/2023.02.28.530532

**Authors:** Kejun Ying, Alexander Tyshkovskiy, Alibek Moldakozhayev, Hanchen Wang, Cecília G. De Magalhães, Sharif Iqbal, Amanda E. Garza, Albina Tskhay, Jesse R. Poganik, Kexin Huang, Yuanhao Qu, Dmitrii Glubokov, Cheng Jin, Donghyun Lee, Hanna Liu, Carolina Leote, Alexandre Trapp, Lucas Paulo de Lima Camillo, Csaba Kerepesi, Mahdi Moqri, Odin Zhang, Kaiyi Jiang, Fedor Galkin, Alex Zhavoronkov, Jeremy M. Van Raamsdonk, Mengdi Wang, Le Cong, Aviv Regev, Jure Leskovec, Tony Wyss-Coray, Vadim N. Gladyshev

## Abstract

Decades of publicly available molecular studies have generated millions of samples testing diverse interventions, yet these datasets were rarely analyzed for their effects on aging. Aging clocks now enable biological age estimation and life outcome prediction from molecular data, creating an opportunity to systematically mine this untapped resource. We developed ClockBase Agent, a publicly accessible platform that reanalyzes millions of human and mouse methylation and RNA-seq samples by integrating them with over 40 aging clock predictions. ClockBase Agent employs specialized AI agents that autonomously generate aging-focused hypotheses, evaluate intervention effects on biological age, conduct literature reviews, and produce scientific reports across all datasets. Reanalyzing 43,602 intervention-control comparisons through multiple aging biomarkers revealed thousands of age-modifying effects missed by original investigators, including over 500 interventions that significantly reduce biological age (e.g., ouabain, KMO inhibitor, fenofibrate, and NF1 knockout). Large-scale systematic analysis reveals fundamental patterns: significantly more interventions accelerate rather than decelerate aging, disease states predominantly accelerate biological age, and loss-of-function genetic approaches systematically outperform gain-of-function strategies in decelerating aging. As validation, we show that identified interventions converge on canonical longevity pathways and with strong concordance to independent lifespan databases. We further experimentally validated ouabain, a top-scoring AI-identified candidate, demonstrating reduced frailty progression, decreased neuroinflammation, and improved cardiac function in aged mice. ClockBase Agent establishes a paradigm where specialized AI agents systematically reanalyze all prior research to identify age-modifying interventions autonomously, transforming how we extract biological insights from existing data to advance human healthspan and longevity.

## Introduction

Biological aging represents the single greatest risk factor for chronic diseases and mortality, yet systematic approaches to identify interventions that modify aging trajectories remain limited^1–3^. Recent advances in machine learning have produced increasingly sophisticated aging clocks that can accurately predict chronological age and capture biological aging processes that correlate with health outcomes and mortality^4,5^. DNA methylation clocks have evolved from first generation models trained solely on chronological age (Horvath 2013, Hannum 2013) to second generation clocks incorporating phenotypic information and mortality risk (PhenoAge, GrimAge, GrimAge2)^6–10^, pace-of-aging measures that capture dynamic aging rates (DunedinPOAm38, DunedinPACE), and most recently causality-enriched models that distinguish causal from correlational aging signals (CausAge, DamAge, AdaptAge)^11^, as well as a pan-mammalian species clock^12^. Similarly, transcriptomic clocks leverage gene expression data to assess chronological age and mortality rate, offering insights into mechanisms of aging and interventions^13–16^. In addition, module-specific transcriptomic clocks have been developed to predict biological age at the level of individual functional modules, providing a higher-resolution assessment of aging biomarkers^14^.

Despite these advances, most clocks have been applied within individual studies under narrow hypotheses. The Gene Expression Omnibus (GEO) represents decades of collective research effort, containing millions of human and mouse molecular profiles from thousands of independent studies testing diverse interventions, genetic perturbations, disease models, and environmental conditions^17^. However, these datasets were generated to address specific research questions unrelated to aging. The original investigators examined their samples for disease mechanisms, drug responses, or tissue-specific processes, but virtually never assessed whether their interventions modified biological age. This represents a massive missed opportunity: a compendium-scale resource where the aging implications of most experiments remain unexplored. The field lacks any systematic, standardized reanalysis of this population-scale archive through the lens of biological age, leaving countless potential age-modifying interventions hidden in plain sight across all past research.

The emergence of large language models and AI agents presents an unprecedented opportunity to systematically reanalyze all biological data at compendium scale^18–20^. While traditional bioinformatic approaches require predefined hypotheses and manual analysis workflows, AI agents can autonomously generate hypotheses, execute complex analytical pipelines, synthesize findings across multiple data sources, and identify patterns that human researchers might overlook. This capability becomes particularly powerful when combined with molecular aging clock algorithms to reanalyze every public dataset. By applying standardized aging biomarkers across all previous studies, we can extract aging-relevant insights from experiments never designed to test aging interventions, effectively transforming the entire historical record of molecular research into an aging intervention discovery engine.

Here we present ClockBase Agent, the first comprehensive platform that systematically reanalyzes all publicly available molecular aging data by integrating approximately 2 million human and mouse samples from Gene Expression Omnibus studies with over 40 aging clock predictions (**Figure 1a**). ClockBase Agent employs an AI agent framework that autonomously processes metadata, generates aging-focused hypotheses, performs standardized statistical analyses, conducts literature reviews, and produces scientific reports across all datasets (**Figure 1b**). Analysis of 43,602 intervention-control comparisons from 13,211 mouse RNA-seq studies revealed thousands of age-modifying effects missed by original investigators. We experimentally validated ouabain, a top-scoring AI-identified candidate not previously studied for anti-aging effects. Treatment with ouabain significantly reduced frailty progression, improved cardiac function, and decreased neuroinflammation in aged mice, demonstrating how systematic reanalysis can reveal actionable health-promoting interventions. The entire resource is freely available.

**Figure 1.**
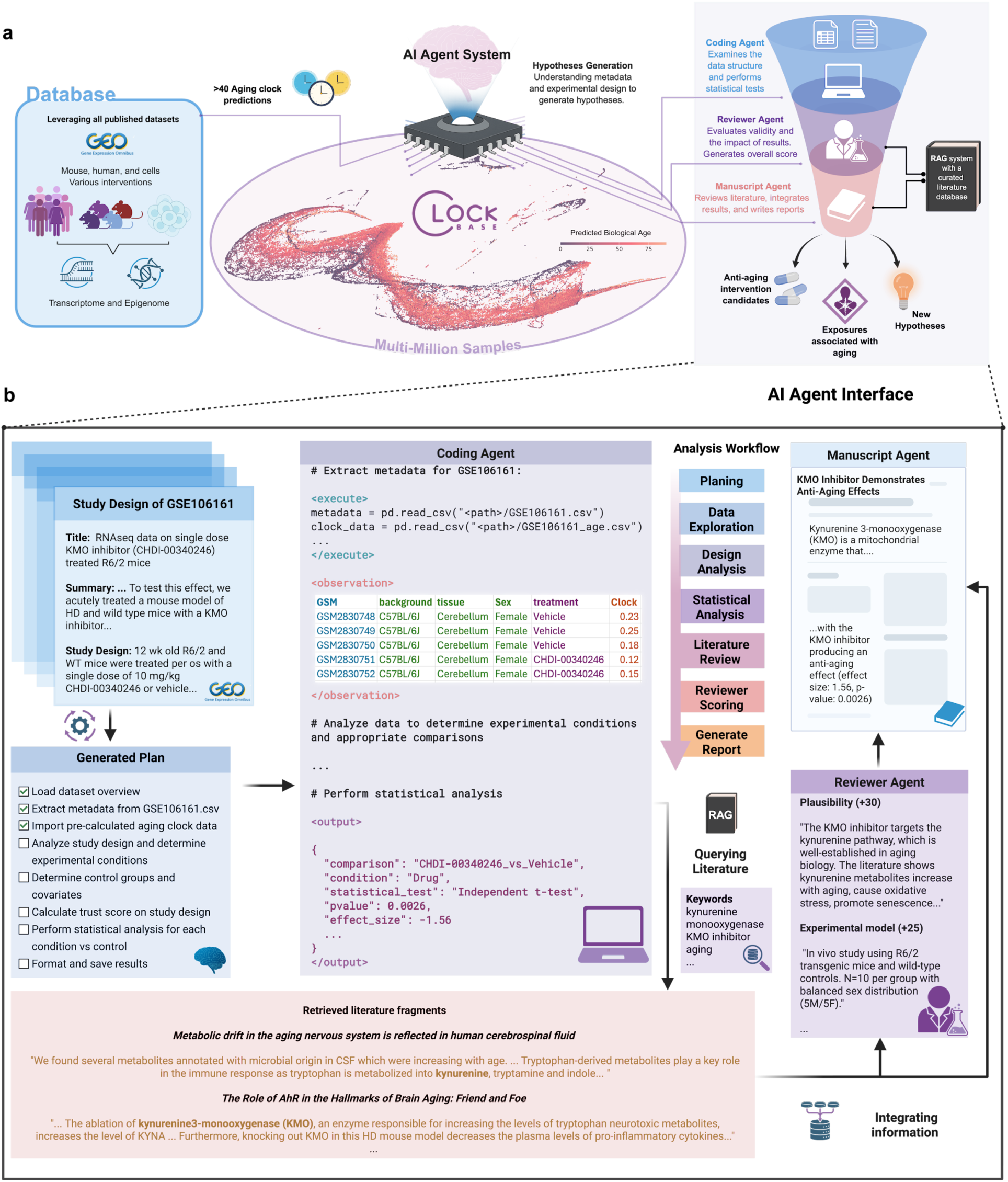
ClockBase Agent platform architecture and workflow. **a.** Schematic diagram showing the system architecture with three main components. The left panel displays the data sources. The center panel shows a UMAP plot with dots colored on a gradient scale representing predicted biological ages. The right panel shows a hierarchical flowchart with three AI agents. **b.** A detailed example of AI agent workflow with (GSE106161). Agent outputs are truncated for illustration purposes.

## Results

ClockBase Agent integrates three core components to enable systematic discovery of age-modifying interventions from all previous molecular research (**Figure 1a**). First, we constructed a comprehensive data integration layer that harmonizes all publicly available human and mouse methylation and RNA-seq samples with standardized aging clock predictions. Second, we implemented a multi-agent AI system where specialized agents autonomously process each dataset through a structured workflow: parsing experimental metadata, generating aging-focused hypotheses, selecting appropriate statistical methods, conducting literature reviews, and producing scientific reports with composite scoring to prioritize interventions (**Figure 1b**). Third, we developed an interactive web platform that provides queryable access to all biological age predictions and analysis results. This integrated architecture enables automated reanalysis of thousands of studies through multiple aging biomarkers.

### A comprehensive atlas of biological age across 2 million samples

To construct a population-scale resource for aging research, we systematically processed human and mouse DNA methylation and RNA-seq samples deposited in the GEO before 2025. ClockBase integrates molecular aging data at unprecedented scale, encompassing 2,048,729 samples processed from publicly available datasets (**Figure 2**). The database comprises 230,516 human DNA methylation samples, 1,749 mouse DNA methylation samples, 852,381 human RNA-seq samples, and 964,083 mouse RNA sequencing samples. For each dataset, we curated experimental metadata and harmonized data formats.

**Figure 2.**
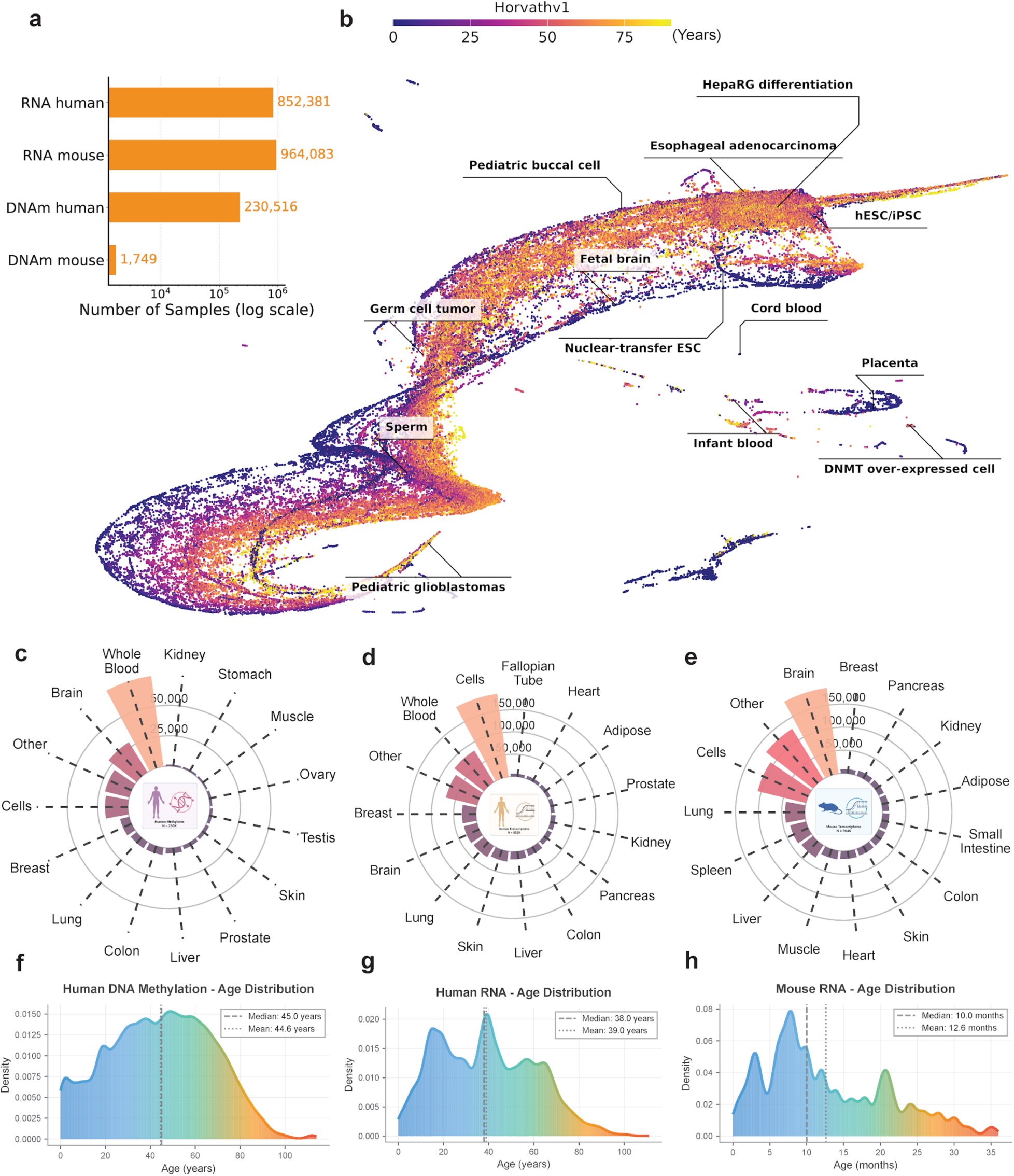
Comprehensive biological age atlas across human and mouse molecular profiles. **a.** The distribution of 2,048,729 samples across four data types: mouse RNA-seq (n=964,083), human RNA-seq (n=852,381), DNA methylation (n=230,516), and mouse DNA methylation (n=1,749). The X axis are on log scale for illustration purposes. **b.** Two-dimensional UMAP plot of human DNA methylation samples (n=230,516). The x-axis represents UMAP dimension 1 (arbitrary units), and the y-axis represents UMAP dimension 2 (arbitrary units). Each dot represents one sample, colored according to the Horvath multi-tissue clock prediction on a continuous scale as shown in the color bar labeled. **c-e.** Circular bar plots showing tissue distribution for human DNA methylation (c), human RNA (d), and mouse RNA (e) datasets. The top 15 tissues ranked by sample count are displayed in descending order. Bar height represents sample count with radial tick marks indicating scale. **f-h.** Age distribution density plots for human DNA methylation (f, n=98,917), human RNA (g, n=87,743), and mouse RNA (h, n=224,009). Curves show kernel density estimates with gradient coloring from cool (blue, younger) to warm (red, older) representing the age spectrum. Shaded areas under curves represent density. Dashed lines indicate median age and dotted lines indicate mean age.

We implemented over 40 aging clock models spanning epigenetic and transcriptomic approaches. These included 39 human DNA methylation clocks capturing chronological age, mortality risk, pace-of-aging, and causality-enriched signals; 4 mouse methylation clocks; and 4 transcriptomic clocks for both chronological age and mortality prediction in mouse and human^6–12,14^ (Methods). The resulting atlas provides biological age estimates with searchable annotations describing disease status, environmental exposures, genetic perturbations, and pharmacological treatments (**Figure 1**). All biological age predictions and analysis results are accessible through an interactive web platform that enables online statistical analysis (e.g., group comparisons, correlation analyses, and accuracy assessments) across all datasets (**Extended Data Figure 1**).

Cross-clock analyses revealed both shared and distinct signals across the comprehensive panel of biomarkers. We observed that correlations among first-generation chronological age predictors (Horvath, Hannum, Lin) remained strong across samples, while second-generation phenotype- and mortality-focused clocks (GrimAge, GrimAge2, PhenoAge) formed distinct clusters reflecting their training on healthspan outcomes (**Extended Data Figure 2a,b**). The Uniform Manifold Approximation and Projection (UMAP) embedding of human DNAm samples reveals distinct clustering patterns that align with biological age. Samples group into clusters that range from biologically young to old states, with infant tissue, placenta, and cell lines with disrupted epigenetic enzymes forming discrete clusters (**Figure 2b**). These features establish ClockBase as the most comprehensive public atlas of biological age to date and lay the foundation for automated discovery of age-modifying conditions at scale.

### AI agent system identifies age-modifying interventions

We deployed the ClockBase agent framework to assess intervention effects on biological age across mouse RNA-seq datasets. After quality control and outlier filtering (Methods), we analyzed 43,529 intervention-control comparisons from 13,211 datasets, encompassing genetic perturbations (20,033), pharmacological treatments (7,933), environmental exposures (4,459), and disease models (3,416). Among these, 5,756 (13.2%) showed statistically significant age-modifying effects (FDR < 0.05) (**Figure 3a-e**, **Figure 4a**, **Table 1**-**2**). We show that the composite score ranking gives meaningful prioritization for the identified age-modifying interventions (**Figure 3b-e, Methods**).

**Figure 3.**
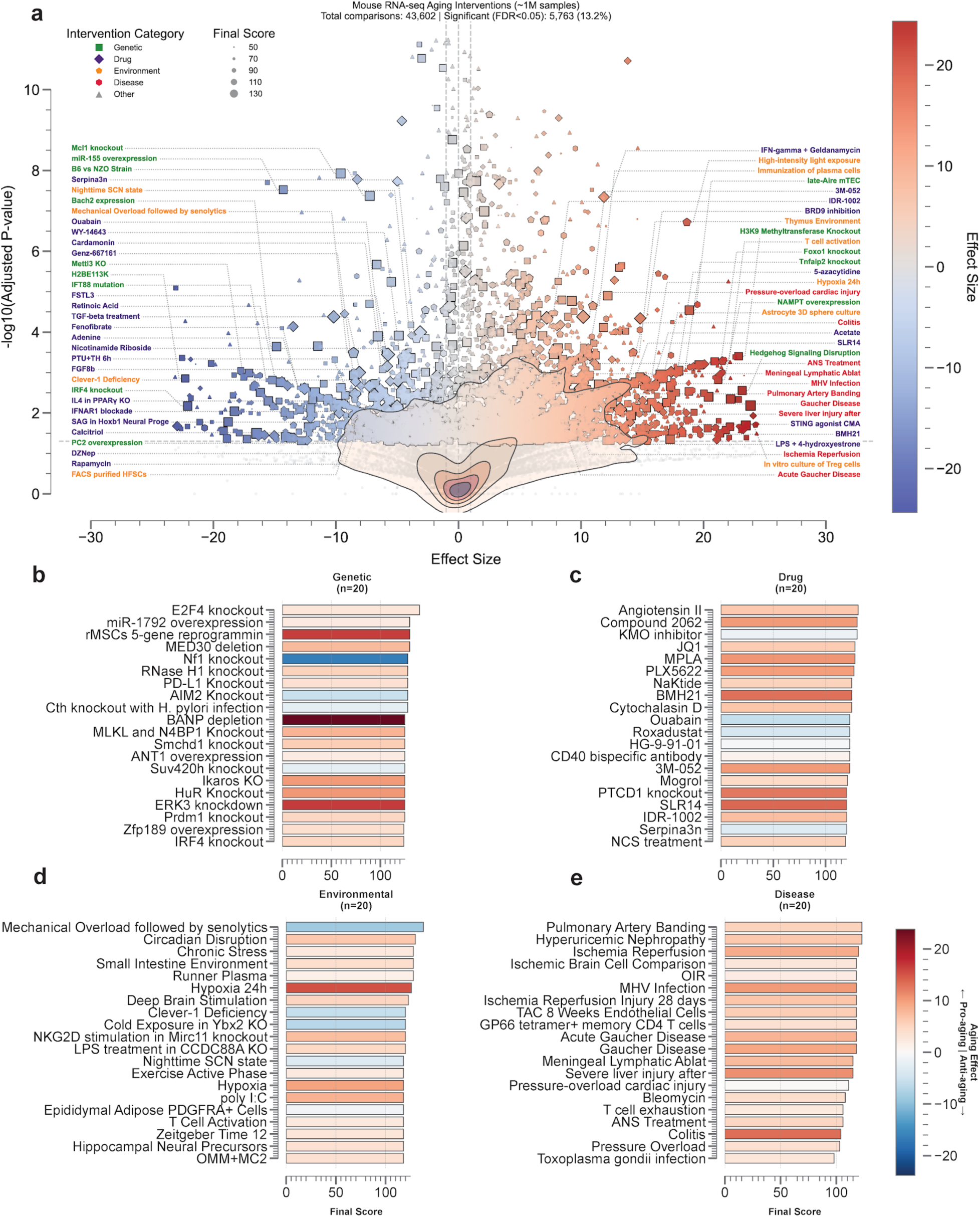
AI agent-identified age-modifying interventions in mouse RNA-seq datasets. **a.** Scatter plot displaying the relationship between effect size and statistical significance for 43,602 age-matched intervention-control comparisons from 13,211 mouse RNA-seq datasets. The x-axis shows log2 effect size (β coefficient) based on mouse RNA mortality clock, where negative values indicate biological age reduction and positive values indicate age acceleration. The y-axis shows -log10(BH-adjusted p-value), with higher values representing greater statistical significance. A horizontal dashed gray line indicates the significance threshold (FDR=0.05). Each point represents a single intervention comparison, with point size proportional to the composite Final Score. Points are colored by intervention category: red for Genetic perturbations (n=20,033), blue for Drug treatments (n=7,933), orange for Environmental exposures (n=4,459), purple for Disease models (n=3,416), and gray for Other interventions (n=7,688). Text annotations label specific interventions at extreme positions on the plot. The legend in the upper left shows intervention categories with total sample sizes and the percentage showing significant effects (p.adjusted<0.05). Background contour shading indicates the density of data points, with darker regions representing higher concentration of interventions. **b-e.** Horizontal bar chart displaying the top 20 genetic interventions ranked by Final Score (all with adjusted p-value<0.05). The x-axis shows Final Score on a 0-150 scale. Bars are colored blue for interventions that reduce biological age (negative β) and red for those that accelerate aging (positive β). Each bar is labeled with the gene name and color represents the effect size.

**Figure 4.**
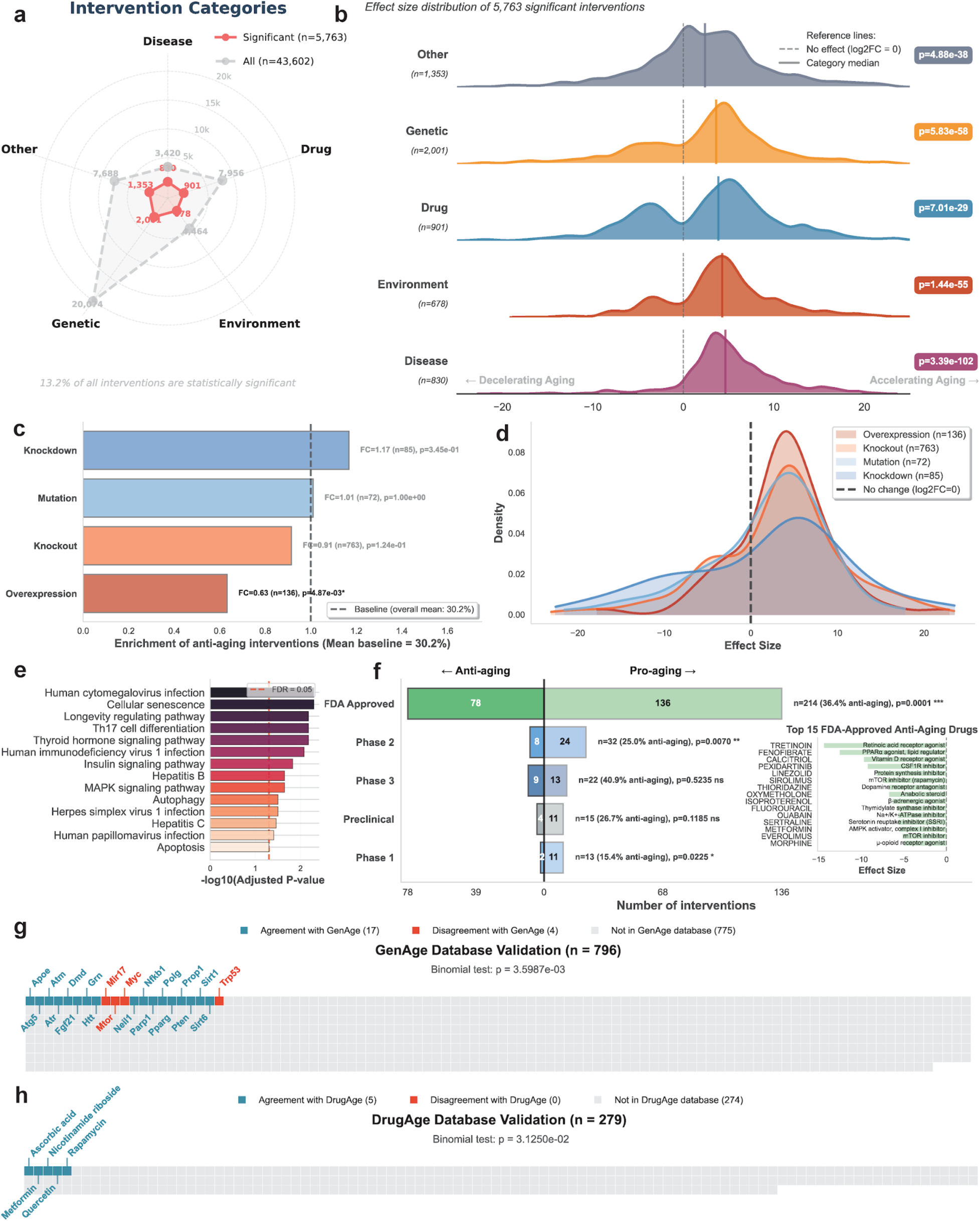
Systematic analysis of intervention categories and biological validation. **a.** Radar plot with five axes representing intervention categories. The gray-filled polygon shows the total number of interventions in each category on a radial scale from 0 to 20,000. The red overlay polygon indicates the proportion of interventions with p.adjusted<0.05. Categories are labeled clockwise from top: Other (n=7,688), Drug (n=7,933), Environment (n=4,459), Disease (n=3,416), and Genetic (n=20,033). Percentages adjacent to each axis indicate the proportion of significant interventions (p.adjusted<0.05) within each category. **b.** Ridge density plot showing the distribution of effect sizes for 5,763 significant interventions (p.adjusted<0.05). The y-axis lists intervention categories ordered by median effect size. The x-axis shows log2 effect size (β coefficient) ranging from -20 to +20. Each ridge represents the kernel density estimate for that category, with ridge height indicating the relative frequency of interventions at each effect size. A vertical gray line at x=0 marks no effect. Ridges are colored by category: gray (Other, n=1,383), orange (Genetic, n=2,001), blue (Drug, n=901), red (Environment, n=678), and purple (Disease, n=830). Adjusted p–values from category enrichment analysis are shown on the right margin. Text labels indicate “Decelerating Aging” (negative values) and “Accelerating Aging” (positive values). **c.** Horizontal bar plot showing fold change enrichment of anti-aging interventions for each genetic intervention type relative to the overall baseline. Knockdown (n=85), knockout (n=763), overexpression (n=136), and mutation (n=72) are shown with bars. Sample sizes and p-values from binomial tests are indicated. **d.** Overlapping density distributions of effect sizes for each genetic intervention type. **e.** Horizontal bar plot of KEGG pathway enrichment analysis results. The y-axis lists the top enriched pathways, and the x-axis shows -log10(adjusted p-value). Bars are colored by enrichment significance on a gradient scale. A vertical dashed red line indicates the FDR=0.05 significance threshold. **f.** Stacked horizontal bar plot showing the distribution of FDA-approved drug interventions by anti-aging (left, darker) and pro-aging (right, lighter) effects. Each bar represents a clinical development stage (FDA Approved, Phase 3, Phase 2, Phase 1, Preclinical) with sample sizes and proportions indicated. An inset panel shows the top 15 FDA-approved anti-aging drugs ranked by effect size, with drug names and mechanisms of action labeled. Statistical significance from binomial tests comparing to 50:50 baseline is shown. **g-h.** Waffle chart validating ClockBase findings against the GenAge and DrugAge database, with blue segment for interventions showing agreement with the database, red segment for disagreement, and gray segment for interventions not present in the database. The overlapped genes are annotated on the chart. The one-sided binomial test P-value is displayed above the chart.

**Table 1:** Top 20 pro-aging interventions identified by AI agents. .

| GSE ID | Intervention | Category | Effect Size | FDR | Final Score | P-value |
| --- | --- | --- | --- | --- | --- | --- |
| GSE109684 | E2F4 knockout | Genetic | 2.96 | 0.049 | 140 | $6.9 \times 10^{-3}$ |
| GSE143809 | Angiotensin II | Drug | 6.35 | 0.024 | 131 | $2.4 \times 10^{-3}$ |
| GSE112232 | rMSCs 5-gene reprogramming | Genetic | 16.53 | $1.1 \times 10^{-3}$ | 130 | $3.7 \times 10^{-5}$ |
| GSE150171 | Compound 2062 | Drug | 10.07 | $5.7 \times 10^{-3}$ | 130 | $3.2 \times 10^{-4}$ |
| GSE181534 | MED30 deletion | Genetic | 7.60 | 0.015 | 130 | $1.2 \times 10^{-3}$ |
| GSE165199 | Circadian disruption | Environment | 6.49 | 0.033 | 130 | $3.8 \times 10^{-3}$ |
| GSE140525 | miR-17~92 overexpression | Genetic | 2.20 | $9.6 \times 10^{-3}$ | 130 | $6.5 \times 10^{-4}$ |
| GSE111730 | MPLA | Drug | 10.44 | 0.014 | 128 | $1.1 \times 10^{-3}$ |
| GSE156126 | JQ1 | Drug | 5.97 | $6.9 \times 10^{-3}$ | 128 | $4.1 \times 10^{-4}$ |
| GSE199138 | RNase H1 knockout | Genetic | 5.43 | 0.030 | 128 | $3.4 \times 10^{-3}$ |
| GSE164401 | Runner plasma | Environment | 1.37 | $5.0 \times 10^{-6}$ | 128 | $6.4 \times 10^{-8}$ |
| GSE185070 | Small intestine environment | Environment | 4.00 | 0.038 | 128 | $4.7 \times 10^{-3}$ |
| GSE148149 | PD-L1 knockout | Genetic | 3.68 | 0.035 | 128 | $4.2 \times 10^{-3}$ |
| GSE138670 | Chronic stress | Environment | 2.54 | 0.025 | 128 | $2.5 \times 10^{-3}$ |
| GSE160660 | PLX5622 | Drug | 10.00 | $2.4 \times 10^{-3}$ | 127 | $1.0 \times 10^{-4}$ |
| GSE196811 | Hypoxia 24h | Environment | 15.23 | $1.5 \times 10^{-3}$ | 126 | $5.1 \times 10^{-5}$ |
| GSE155598 | BANP depletion | Genetic | 23.83 | $6.5 \times 10^{-3}$ | 125 | $3.8 \times 10^{-4}$ |
| GSE142424 | ERK3 knockdown | Genetic | 16.65 | $1.3 \times 10^{-3}$ | 125 | $4.5 \times 10^{-5}$ |
| GSE185424 | BMH21 | Drug | 13.19 | $8.1 \times 10^{-3}$ | 125 | $5.1 \times 10^{-4}$ |
| GSE137323 | HuR knockout | Genetic | 10.29 | $4.5 \times 10^{-3}$ | 125 | $2.3 \times 10^{-4}$ |

**Table 2:** Top 20 anti-aging interventions identified by AI agents.

| GSE ID | Intervention | Category | Effect Size | FDR | Final Score | P-value |
| --- | --- | --- | --- | --- | --- | --- |
| GSE195707 | Mechanical overload followed by senolytics treatment | Environment | -8.57 | $3.3 \times 10^{-4}$ | 138 | $7.7 \times 10^{-6}$ |
| GSE106161 | KMO inhibitor | Drug | -1.56 | 0.025 | 130 | $2.6 \times 10^{-3}$ |
| GSE159021 | Nf1 knockout | Genetic | -16.48 | 0.032 | 128 | $3.7 \times 10^{-3}$ |
| GSE184479 | AIM2 knockout | Genetic | -5.48 | 0.011 | 128 | $7.9 \times 10^{-4}$ |
| GSE158817 | Cth knockout with H. pylori infection | Genetic | -2.40 | 0.042 | 128 | $5.4 \times 10^{-3}$ |
| GSE117482 | Suv420h knockout | Genetic | -2.39 | 0.042 | 125 | $5.5 \times 10^{-3}$ |
| GSE81424 | Nkx3.1 knockout | Genetic | -6.58 | $1.3 \times 10^{-4}$ | 123 | $2.6 \times 10^{-6}$ |
| GSE122080 | Ouabain | Drug | -5.74 | $3.6 \times 10^{-4}$ | 123 | $8.6 \times 10^{-6}$ |
| GSE150669 | Roxadustat | Drug | -4.27 | 0.028 | 123 | $3.1 \times 10^{-3}$ |
| GSE163972 | HG-9-91-01 | Drug | -0.73 | $2.3 \times 10^{-5}$ | 123 | $3.6 \times 10^{-7}$ |
| GSE137096 | VEC-Cre endothelial mutagenesis | Genetic | -4.50 | $5.6 \times 10^{-4}$ | 122 | $1.5 \times 10^{-5}$ |
| GSE102227 | Mcl1 knockout | Genetic | -9.62 | $1.2 \times 10^{-8}$ | 120 | $1.0 \times 10^{-10}$ |
| GSE159795 | B6 vs NZO strain comparison | Genetic | -7.24 | $4.2 \times 10^{-8}$ | 120 | $3.9 \times 10^{-10}$ |
| GSE183055 | IPr subunit expression | Genetic | -11.72 | $3.1 \times 10^{-3}$ | 120 | $1.4 \times 10^{-4}$ |
| GSE167301 | SETBP1 dominant-negative | Genetic | -7.76 | 0.012 | 120 | $9.3 \times 10^{-4}$ |
| GSE148730 | Clever-1 deficiency | Environment | -5.69 | $5.1 \times 10^{-3}$ | 120 | $2.7 \times 10^{-4}$ |
| GSE86590 | Cold exposure in Ybx2 KO | Environment | -6.82 | 0.014 | 120 | $1.1 \times 10^{-3}$ |
| GSE128145 | CtBP2 mutant | Genetic | -3.90 | $4.4 \times 10^{-4}$ | 120 | $1.1 \times 10^{-5}$ |
| GSE148794 | Serpina3n | Drug | -3.12 | $1.0 \times 10^{-4}$ | 120 | $1.9 \times 10^{-6}$ |
| GSE176174 | BHLHE40 knockout | Genetic | -7.43 | 0.034 | 120 | $4.1 \times 10^{-3}$ |

Analysis of 20,033 genetic perturbations revealed that 1,996 (10.0%) significantly altered tAge (Figure 3b). The most striking genetic interventions that reduced biological age included IRF4 knockout, a key transcription factor in immune cell differentiation (=−22.10, P = 3.97 × 10^−4^, score = 115), and Mettl3 knockout, which disrupts m6A RNA methylation (=−17.10, P = 3.19 × 10^−5^, score = 105). Notably, both Bach2 expression, a transcriptional repressor that maintains T cell quiescence (=−14.85, P = 5.00 × 10^−6^, score = 115), and miR-155 overexpression, despite its pro-inflammatory role (=−14.29, P = 2.69 × 10^−10^, score = 108), paradoxically reduced biological age. Conversely, disruption of hedgehog signaling, critical for tissue homeostasis (=23.15, P = 1.21 × 10^−4^, score = 112), and knockout of H3K9 methyltransferases, essential for heterochromatin maintenance (=22.87, P = 9.73 × 10^−6^, score = 105), substantially accelerated aging. The composite scoring analysis prioritized interventions combining strong effect sizes with biological plausibility and experimental rigor.

Among the 7,933 pharmacological interventions, 900 (11.3%) produced a statistically significant effect on transcriptomic age (tAge) as assessed by the mouse mortality predictor^14^. The most notable age-reducing drugs (**Figure 3c**) include ouabain, a cardiac glycoside with established senolytic properties (=−5.74, P = 8.59 × 10^−6^, score = 123), the mTOR inhibitor rapamycin (=−7.50, P = 9.83 × 10^−4^, score = 105), and fenofibrate, a PPARα agonist used clinically for dyslipidemia (=−12.57, P = 1.04 × 10^−4^, score = 111). Among immune modulators, Serpina3n, a serine protease inhibitor, showed strong age-reducing effects (=−3.12, P = 1.94 × 10^−6^, score = 120). Conversely, interventions that accelerated aging included BMH21, a nucleolar stress inducer that disrupts ribosomal RNA synthesis (=13.19, P = 5.15 × 10^−4^, score = 125), and the innate immune activator 3M-052, a TLR7/8 agonist (=10.18, P = 7.02 × 10^−7^, score = 123).

Environmental and disease perturbations showed distinct patterns across 4,459 environmental exposures and 3,416 disease models. Among environmental factors (**Figure 3d-e**), mechanical overload, representing mechanical damage coupled with senolytic treatment (=−8.57, P = 7.67 × 10^−6^, score = 138), significantly reduced biological age. In contrast, high-intensity light exposure on early-stage embryos (=18.64, P = 2.00 × 10^−9^, score = 111) and hypoxia (=15.23, P = 5.05 × 10^−5^, score = 126) accelerated aging. In the case of diseases, as expected, pathological conditions predominantly accelerated aging, including ischemia-reperfusion injury (=8.92, P = 5.95 × 10^−4^, score = 120), viral infection (MHV; =10.18, P = 2.63 × 10^−4^, score = 118), and metabolic disorders such as Gaucher disease (=9.28, P = 2.78 × 10^−4^, score = 118). The data revealed a clear dichotomy: while most disease states accelerate biological aging, certain therapeutic interventions and adaptive stressors were predicted to reverse mortality-associated trajectories.

### Systematic characterization of age-modifying interventions

Our analysis revealed category-specific patterns (**Figure 4a-b**). Disease states showed the highest significance rate (24.3%), followed by environmental exposures (15.2%), drug treatments (11.3%), and genetic interventions (10.0%). Effect sizes ranged from −23.12 to 24.35 (median 0.33), with a positive skew indicating that interventions more commonly accelerate than decelerate aging biomarkers. The agent’s composite scoring system (median 85) prioritized interventions based on study design and biological relevance, with 33.0% exceeding 100 points.

To determine whether specific genetic intervention strategies differ in their anti-aging potential, we classified 1,056 significant genetic interventions by type: knockdown (n=85), knockout (n=763), overexpression (n=136), and mutation (n=72) (**Figure 4c**, **Methods**). Overexpression yielded significantly fewer anti-aging interventions (19.1%) compared to the overall baseline of 30.2% (p=0.005, fold change=0.63). In contrast, both loss-of-function approaches showed significant enrichment. Knockdown interventions demonstrated 1.85-fold higher anti-aging proportions than overexpression (35.3% vs 19.1%, p=0.011, odds ratio=2.31), while knockout interventions showed 1.45-fold enrichment (27.7% vs 19.1%, p=0.044, odds ratio=1.62). The distribution of effect sizes across intervention types (**Figure 4d**) revealed that overexpression interventions were more likely to accelerate aging, while knockdown and knockout interventions showed broader distributions extending into the anti-aging range. These findings suggest that overexpression interventions are more likely to be damaging rather than beneficial, consistent with the established notion that gene overexpression can disrupt cellular homeostasis. This pattern indicates that overexpression-based interventions require particularly careful design and validation, while loss-of-function approaches may offer a more favorable risk-benefit profile for aging intervention development.

To validate biological plausibility, we performed pathway enrichment analysis on the significant genetic interventions using Over-Representation Analysis (ORA) with KEGG pathway databases (**Figure 4e**). Among the validated genes, we identified significant enrichment in established aging-related pathways (FDR < 0.05). The top-ranking pathways included cellular senescence, longevity regulating pathway, Th17 cell differentiation, and human cytomegalovirus infection. Additional enriched pathways encompassed thyroid hormone signaling, human immunodeficiency virus 1 infection, insulin signaling pathway, hepatitis B, MAPK signaling pathway, autophagy, herpes simplex virus 1 infection, hepatitis C, human papillomavirus infection, and apoptosis. This significant convergence on canonical aging mechanisms validates that aging clock perturbations reflect genuine biological aging processes rather than technical artifacts.

To demonstrate clinical relevance, we classified 214 OpenTargets-matched significant drug interventions by clinical development stage (**Figure 4f**)^21^. Clinical development stage significantly associated with enrichment of anti-aging interventions (χ²=10.61, p=0.031). Among FDA-approved drugs, we identified 78 compounds showing anti-aging effects (36.4% anti-aging rate) and 136 showing pro-aging effects, with significantly more pro-aging drugs than anti-aging drugs (n=214, p=0.0001). The top 15 FDA-approved anti-aging drugs identified include tretinoin, fenofibrate, calcitriol, pexidartinib, linezolid, as well as, well known anti-aging drugs such as sirolimus and metformin.

To benchmark against curated longevity knowledge, we compared all significant and high-confidence gene perturbations (n = 796; FDR < 0.05 and final score > 80) with curated entries in the GenAge database^22^. A total of 21 unique genes overlapped with GenAge annotations (**Figure 4g**). Among these, 17 interventions (81.0%) showed directional agreement (e.g., *Grn*, *Pten*, *Neil1*, and *Apoe*). A one-sided binomial test yielded *P* = 3.60 × 10^-3^, indicating significant enrichment of concordant longevity effects among matched interventions. Four interventions showed directional mismatches. These discrepancies are partially attributable to the context-specific nature of genetic perturbations. For example, although *Mtor* is classified as an anti-longevity gene based on systemic inhibition studies, complete or lineage-restricted knockout can produce detrimental metabolic and physiological effects that increase biological age. We performed a similar validation for pharmacological perturbations (**Figure 4h**). Among 279 high-confidence drug interventions, only 5 treatments (1.8 percent) were present in the DrugAge database. All 5 matched compounds showed perfect directional concordance with DrugAge annotations (100%, binomial test *P* = 0.031). These included rapamycin, nicotinamide riboside, quercetin, ascorbic acid, and metformin, each classified as pro-longevity or anti-longevity in agreement with the database. The remaining 775 genetic interventions (97.4%) and 274 drug interventions (98.2%) were not previously linked to lifespan in GenAge and DrugAge, indicating that the agent framework reveals a substantial set of new intervention candidates for aging research.

### Agent operation patterns demonstrate sophisticated analytical capabilities

Analysis of 17,695 agent execution traces across the complete mouse RNA-seq corpus revealed systematic decision-making patterns that closely mirror expert bioinformatic workflows (**Figure 1b**, **Figure 5a-c**). The agent autonomously progressed through standardized analytical phases: metadata parsing and quality assessment, experimental design evaluation (identifying treatment groups, controls, and covariates), statistical method selection (choosing between t-tests, Welch’s tests, linear models, or z-tests based on sample size and variance homogeneity). The systematic workflow is demonstrated through operation traces across datasets (**Figure 5b**) and state transition patterns (**Figure 5c**).

**Figure 5.**
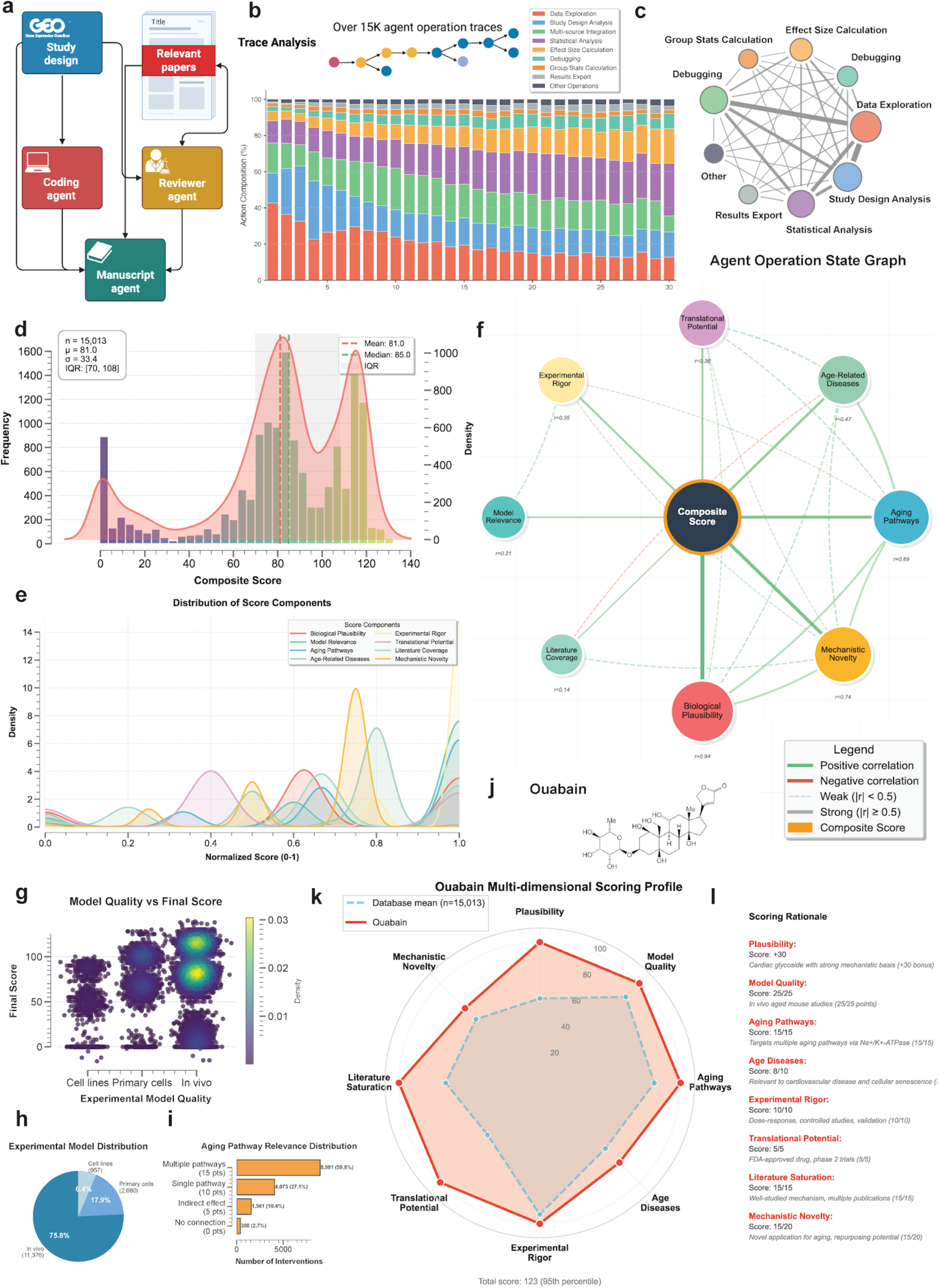
AI agent operational workflow and multi-dimensional scoring system. **a.** Schematic plot illustrating the information flow among AI agent system. **b.** Stacked bar chart showing operation traces across 17,695 scored comparisons. The x-axis represents steps, and the y-axis shows operation percentage. Each bar consists of colored segments representing different actions. **c.** Network graph displaying agent state transitions. Circular nodes represent different analytical states, with the central Composite Score node. Peripheral nodes include Group State Calculation, Debugging, Data Exploration, Study Design Analysis, Results Export, and Statistical Analysis. Edge thickness is proportional to transition frequency (n=17,695 total traces). **d.** Histogram showing the distribution of composite Final Scores. The x-axis shows score values ranging from - 20 to 140, and the y-axis shows frequency. Vertical dashed lines mark quartiles. **e.** Density plot showing the distribution of each score component across 17,695 scored comparisons. The color represents a different sub-score and the x axis shows the normalized score (scaling from 0 to 1). **f.** Correlation network showing relationships between scoring components. Circular nodes are sized proportionally to mean score and colored by variance (light to dark blue gradient). Edges represent Pearson correlations with |r|>0.3, with blue edges for positive correlations and red for negative. Edge thickness scales with correlation strength. **g.** Scatter plot examining the relationship between Model Quality and Final Score. The x-axis shows Model Quality categories, and the y-axis shows Final Score. Points are colored by kernel density. **h.** Pie chart showing the distribution of experimental model types among scored interventions. **i.** Bar plot showing the distribution of aging pathway relevance score across interventions. **j.** The molecular graph of ouabain. **k.** Overlaid radar plots comparing ouabain’s scoring profile (red polygon) against median scores for all drug interventions (blue polygon). Eight axes represent the same scoring dimensions as panel e. **l.** Table displaying ouabain’s individual scores across eight evaluation dimensions. Each cell shows the score achieved and maximum possible (e.g., “25/25”).

The agent demonstrated consistent analytical rigor across diverse experimental contexts. Analysis traces included systematic verification steps: sample size validation (rejecting analyses with n < 3), outlier detection using interquartile range methods, covariate adjustment when specified in metadata, and false discovery rate correction using the Benjamini-Hochberg procedure. The agent appropriately handled edge cases, including zero-variance groups (applying z-tests for single-sample comparisons), missing metadata fields (inferring from sample naming conventions), and batch effects (incorporating batch as a covariate when detected).

Manual validation of 100 randomly selected analytical traces by two independent doctoral-level bioinformaticians confirmed the agent’s high analytical accuracy across multiple dimensions (**Extended Data Figure 4a-d**). The validation examined eight key criteria: hypothesis formulation, statistical test selection, p-value validity, effect size calculation, comparison validity, analytical completeness, data integration, and protocol adherence (**Extended Data Figure 4a**). Across 1,600 total assessments (800 per evaluator), the agent achieved an overall correctness rate of 99.27% (1488/1499 analyzable criteria, **Extended Data Figure 4b,c**), while 101 cases (6.31%) reflected data-related constraints such as insufficient sample sizes (n < 2 per group) or missing values in the source data. These data-driven limitations were noted but not counted against the agent’s performance, as they stem from experimental design choices or data quality issues beyond the agent’s control. Excluding these data-limited cases, the agent demonstrated 99.27% correctness across 1,499 analyzable assessments (**Extended Data Figure 4d**).

Agent errors fell into three primary categories: failure to identify clear control versus treatment comparisons (4 cases across both evaluators), selection of unrecognized statistical test descriptions (4 cases), and execution of forbidden direct age comparisons such as “young versus old” (3 cases). The most common issue involved hypothesis clarity, where the agent occasionally struggled to articulate explicit control-treatment relationships in studies with complex experimental designs. Direct age comparisons violated the protocol requirement to analyze aging clock effects rather than chronological age differences, representing a conceptual rather than computational error.

To enable the efficient access of AI agents to relevant research papers, we developed a lightweight, cost-effective Retrieval-Augmented Generation (RAG) framework (**Methods**). This system is engineered for robust performance while optimizing for rapid deployment and minimal operational overhead. We indexed over 6,000 aging-related papers from the past five years into a vector database to provide domain-specific context.

The multi-dimensional scoring system employed by the agent integrated eight distinct evaluation criteria to prioritize therapeutic candidates from 15,013 scored comparisons (**Figure 5d-h**, **Methods**, **Extended Data Figure 5**). The scoring algorithm weighted biological plausibility (-50, 0, or +30 points, mean = 6.0), experimental model quality (0-25 points, mean = 21.6), relevance to aging pathways (0-15 points, mean = 12.2), association with age-related diseases (0-10 points, mean = 6.6), experimental rigor (0-10 points, mean = 9.3), translational potential (0-5 points, mean = 2.6), literature saturation (0-15 points, mean = 10.0), and mechanistic novelty (0-20 points, mean = 12.8). The distribution of composite scores (**Figure 5d-f**) ranged from -16 to 140, with a median of 85. The strong relationship between model quality and final score (**Figure 5g**) reflects the database’s emphasis on physiologically relevant systems, with 75.8% of studies using *in vivo* models (**Figure 5h**). High-scoring interventions (≥100) comprised 33.7% of all comparisons and consistently demonstrated superior experimental quality and mechanistic relevance. This systematic prioritization enabled efficient identification of promising candidates, with ouabain achieving a score of 123 among pharmacological interventions suitable for experimental validation (**Figure 5i-k**).

The plausibility filter identified 1,839 interventions (12.3%) exhibiting fundamental biological implausibility that received -50 point penalties, while 6,074 interventions (40.5%) received +30 point bonuses for strong mechanistic rationale.

Aging pathway relevance analysis demonstrated that 8,981 interventions (59.8%) showed direct modulation of multiple established aging mechanisms, while 4,073 (27.1%) affected single pathways. The top-scoring interventions included E2F4 knockout (score = 140), Snf2h knockout (139), and FNDC5 knockout (138). The scoring framework effectively balanced experimental quality with mechanistic innovation, providing a quantitative basis for prioritizing interventions for further investigation.

### Ouabain validation demonstrates therapeutic efficacy in aged mice

Among the compounds predicted to reduce transcriptomic age, we identified ouabain, a cardiac glycoside with established senolytic activity and the ability to improve the immune and physical functions of aged mice^23^. Both chronological and mortality multi-tissue clocks predicted a statistically significant reduction in tAge in 26-month-old C57Bl/6 female mice treated intermittently with ouabain for 13 weeks, compared to age-matched saline controls (**Figure 6a**, **Extended Data Figure 6a**). According to the mortality clock, the top gene contributing to the reduced tAge was *Nrep*, which was significantly upregulated in ouabain-treated mice (**Figure 6b**). Notably, this gene, encoding the neuronal regeneration-related protein, was also among the key contributors to the rejuvenating effect of heterochronic parabiosis in aged mice^14^. Module-specific multi-tissue clocks revealed multiple pathways underlying the anti-aging effects of ouabain, including those linked to inflammation, mRNA splicing, Nrf2 and interferon signaling, translation, lipid metabolism, and oxidative phosphorylation (**Extended Data Figure 6b**). These results suggest that ouabain may exert systemic effects on aged murine livers.

**Figure 6.**
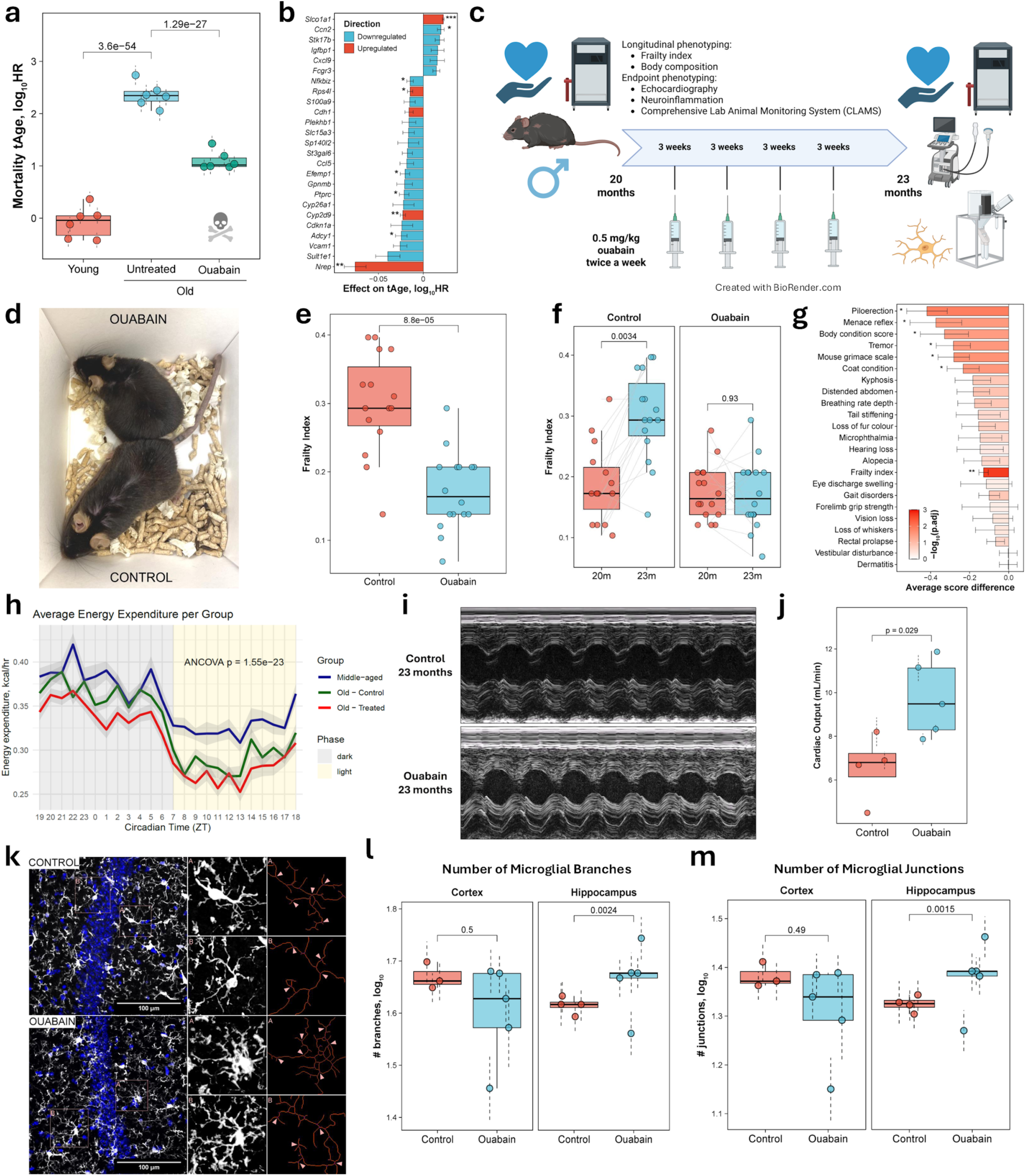
Experimental validation of ouabain’s geroprotective effects in aged mice. **a.** Transcriptomic age (tAge) predicted by rodent multi-tissue clock of expected mortality for young (7-week-old), old (24-month-old) untreated and ouabain-treated C57BL/6 female mice (n=6 per group). Data are mean tAges ± s.d. **b.** Top 25 genes contributing to transcriptomic age increase (positive values) or decrease (negative values) in ouabain-treated mice compared to age-matched controls. The x-axis shows contribution magnitude (calculated as log fold-change multiplied by clock coefficient). Genes up- and downregulated in response to ouabain are colored in red and blue, respectively. **c.** Timeline schematic of the experimental protocol. The list of assessments after 3 months of treatment is denoted in text. **d.** Representative photographs of 23-month-old mice after 3 months of treatment with ouabain or PBS (control injection). Visual differences in posture and coat condition are apparent. **e.** Frailty index (FI) of control (red) and ouabain-treated animals (blue) after 3 months of treatment (n=15-16 per group). Each data point represents an individual mouse. P-value derived with the Wilcoxon rank-sum test is shown in text. **f.** Dynamics of frailty index over 3 months in old control and ouabain-treated animals. FI before (red) and after (blue) treatment is shown for both control (left) and ouabain-treated animals (right). Change in FI over time was assessed for each group with the Wilcoxon signed-rank test. P-values were adjusted for multiple comparisons with the BH method. **g.** Difference in average scores of individual frailty parameters induced by ouabain after 3 months of treatment compared to age-matched control animals. P-values were derived with the Wilcoxon rank-sum test and adjusted for multiple comparisons with the BH method. **h.** Circadian energy expenditure traces over 24 hours (averaged 48-hour data) in middle-aged and old control and ouabain-treated male mice. The x-axis shows time in hours. Light and dark phases are depicted with the background color. Experimental groups are shown with line colors. Shaded bands around lines indicate standard error. **i.** M-mode echocardiography traces. Top and bottom panels show a representative control and ouabain-treated mouse hearts, respectively. Waveforms display cardiac cycles showing systole and diastole. **j.** Average cardiac output measurements for control (red) and ouabain-treated (blue) mice after 3 months of injections (n=4-5 mice per group) assessed with echocardiography. P-value derived with a mixed-effect model is shown in text. Data are mean ± s.e.m. **k.** Representative images of Iba1-labelled microglia (white) and DAPI (blue) in the CA1 region of the hippocampus from control (top row) and ouabain-treated (bottom row) mice. For each group, the left panel displays the full-field image, the middle panel shows a magnified view of the boxed area, and the right panel presents the corresponding segmentation and skeletonization of Iba1⁺ microglia, where pink arrows indicate junctions and red lines indicate branches. **l-m.** Average microglial branch **(l)** and junction **(m)** numbers in cortex (left), and hippocampus (right) of control (red) and ouabain-treated mice (blue) after 3 months of injections (n=4-5 mice per group). The y-axis shows mean log-transformed count across images. P-values were derived with the mixed-effect model and adjusted for multiple comparisons with the BH method. Data are mean ± s.e.m. Boxplots: center line indicates median; boxes, interquartile range; whiskers, ±1.5× IQR. Unless otherwise noted, asterisks denote BH-adjusted p-values. * p.adj < 0.05; ** p.adj < 0.01; *** p.adj < 0.001.

To assess whether ouabain treatment influences frailty and other phenotypic features of aged animals, and whether its anti-aging effects extend across sexes, we subjected 20-month-old C57Bl/6 males to intermittent ouabain treatment for 3 months, following a similar experimental protocol as in the original study (**Figure 6c**). For both the control and ouabain-treated groups, we collected aging-associated phenotypic data, including longitudinal assessment of frailty parameters, weight, and body composition, as well as measurements of cardiac and metabolic functions, and neuroinflammation after 3 months of treatment^24^.

At the end of the experiment, we observed clear morphological and behavioral differences between groups (**Figure 6d**), reflected in a significantly lower frailty index (FI) in ouabain-treated mice (**Figure 6e**). While FI did not differ between groups at baseline, it increased significantly in controls over the 3-month treatment period, whereas ouabain-treated mice showed no such increase in this parameter (**Figure 6f**). The ouabain group mice also showed significant improvements across multiple individual frailty parameters, including reduced piloerection, preserved menace reflex, better body condition, reduced tremor, lower mouse grimace scale, and improved coat condition (**Figure 6g**, **Extended Data Figure 6c**). Notably, none of the frailty parameters indicated adverse health outcomes in the ouabain group compared with saline controls.

To assess the impact of aging and ouabain on metabolic function and feeding behavior, we used the comprehensive lab animal monitoring system (CLAMS) to evaluate old (23-month-old) saline-and ouabain-treated male mice as well as middle-aged (13-month-old) control males. Old mice displayed reduced energy expenditure compared to middle-aged controls (**Figure 6h**), particularly during the light phase in both old groups and during the dark phase in ouabain-treated animals. Pairwise comparisons confirmed significantly lower body weight–adjusted energy expenditure in old controls relative to middle-aged mice during the light phase, with a trend toward reduction during the dark phase (**Extended Data Figure 6d**). Ouabain-treated old mice also showed a marginal decrease in energy expenditure relative to saline-treated old controls during the dark phase.

Similarly, body weight-adjusted CO2 production and O2 consumption were higher in middle-aged mice compared to old saline-treated animals: across both phases for CO2 and during the light phase for O2 (**Extended Data Figure 6e** for CO2; O2 not shown). Ouabain treatment induced a marginal reduction in CO2 production during the dark phase (**Extended Data Figure 6f**), consistent with a slight decrease in energy expenditure. Respiratory exchange ratio (RER) values were comparable across all groups (**Extended Data Figure 6g**), further supporting this interpretation.

Food intake during the dark phase was higher in middle-aged animals compared to old saline-treated mice, but no differences were observed between old groups, suggesting that ouabain’s metabolic effects are not attributable to reduced energy intake (**Extended Data Figure 6h**). Locomotor and ambulatory activity were largely similar across groups (**Extended Data Figure 6i** for locomotor activity; ambulatory activity not shown). Finally, energy balance was modestly reduced in old animals (**Extended Data Figure 6j**), though pairwise differences did not reach statistical significance.

Body composition was assessed by EchoMRI at baseline and after 3 months of treatment. Consistent with food intake, activity and energy balance assessments, neither body mass nor fat-to-lean ratio differed significantly between control and ouabain-treated animals at any timepoint (**Extended Data Figures 6k** **and 6l**). However, both groups displayed significant reductions in body mass and fat-to-lean ratio over the course of treatment.

Using echocardiography, we evaluated cardiac function of 23-month-old males after 3 months of intermittent saline or ouabain treatment (**Figure 6i**). Ouabain-treated mice exhibited significantly higher cardiac output compared to controls at the end of the experiment (**Figure 6j**), consistent with its expected role as a cardiac glycoside^23^. Stroke volume and left ventricular internal diameter at end-systole (LVIDs) and end-diastole (LVIDd) did not differ significantly between groups (**Extended Data Figure 6m**). However, ouabain-treated mice showed a trend toward higher stroke volume and diastolic left ventricular internal dimensions, which may indicate that improved heart relaxation contributes to the observed increase in cardiac output.

To investigate whether ouabain modulates age-associated neuroinflammation, we analyzed microglial morphology in the cortex and hippocampus of 23-month-old male mice (**Figure 6k**). In the hippocampus, microglia from ouabain-treated animals exhibited a greater number of branches and junctions compared to controls (**Figures 6l and 6m**), suggesting higher ramification and lower neuroinflammation in response to treatment^25,26^. By contrast, no differences between groups were detected in the cortex. Maximum branch length and total microglial area were also comparable across groups in both regions (**Extended Data Figure 6n** **and 6o**), although hippocampal microglia of ouabain-treated mice showed a trend toward higher values.

## Methods

### Data acquisition and preprocessing

We systematically downloaded all publicly available mouse and human DNA methylation from the Gene Expression Omnibus (GEO) database deposited before January 2025. The RNA-seq datasets used in this work were downloaded from the ARCHS4 resource, which provides uniformly processed gene and transcript counts from GEO/SRA^27^.

For RNA-seq data preprocessing, we followed the standardized pipeline described in Tyshkovskiy et al.^14^: raw counts underwent soft filtering to remove non-expressed genes (requiring at least 10 reads in at least 20% of samples), followed by RLE (Relative Log Expression) normalization, log-transformation, and scaling of gene expression profiles. For each dataset, we calculated relative gene expression by selecting control samples as a reference group and subtracting their median expression profile from all samples within the same dataset and tissue. This approach effectively reduces batch effects and tissue-specific baseline differences. Missing values were imputed using average expression values precalculated from the training sets of the respective clocks.

DNA methylation arrays were processed using minfi for Illumina 450K/EPIC arrays, and the data structure was standardized according to the Biolearn framework. All processed data were harmonized to ensure consistent gene identifiers and probe annotations across datasets.

### Biological age clock implementation

We implemented over 40 aging clock models through multiple specialized frameworks. For human DNA methylation data, we utilized the Biolearn Python library (version 0.3.0, https://bio-learn.github.io) to calculate 39 epigenetic clocks: (1) chronological age predictors including Horvath multi-tissue^8^, Hannum blood^7^, Lin blood^28^, Horvath skin and blood^29^, and PedBE pediatric buccal^30^; (2) mortality and healthspan-related clocks including PhenoAge^9^, GrimAge^10^, GrimAge2^31^, and Zhang 10-CpG mortality risk^32^; (3) pace-of-aging biomarkers DunedinPOAm38^33^ and DunedinPACE^6^; (4) causality-enriched clocks including Ying CausAge, DamAge, and AdaptAge^11^; and (5) specialized predictors for DNAm telomere length^34^ and gestational age^35^. For mouse methylation, we implemented Petkovich blood^36^, Meer multi-tissue^37^, Thompson multi-tissue^38^, and Wang liver^39^ clocks through Biolearn.

For transcriptomic data, we applied multiple clock types to mouse samples^14^: (1) Mouse multi-tissue elastic net (EN) chronological age models trained on samples from 26 tissues of healthy mice and rats; (2) Mouse multi-tissue EN clocks of expected mortality trained on samples from both control animals and those subjected to lifespan-modulating interventions; (3) Mouse multi-tissue EN clocks of normalized age that predict chronological age normalized by expected maximum lifespan (99.9th percentile) for each experimental condition; and (4) Mouse tissue-specific chronological clocks for liver, kidney, brain, and skeletal muscle. For human samples, we applied the multi-species multi-tissue clocks of chronological age and expected mortality from the same framework, which were trained on combined data from human, mouse, crab-eating macaque, and rat samples to enable robust cross-species age prediction.

### AI agent architecture and implementation

ClockBase uses a multi-agent system built on Anthropic Claude 3.7 Sonnet and OpenAI GPT 4O, tailored for biomedical analysis. The system comprises three agents: (1) an analysis executor, a coding agent that ingests GEO records and supplements and then generates and executes code to select appropriate statistical models based on sample size, distribution, and experimental design; (2) a biological interpreter that contextualizes findings within aging biology literature; and (3) a scorer that evaluates interventions across multiple criteria. The system was deployed on cloud infrastructure with automatic scaling to process datasets in parallel.

We analyzed 15,380 successful agent-study interactions (206,543 code execution blocks) from the ClockBase system to characterize the agent behavior patterns. Agent actions were categorized using semantic code analysis that examined code block purpose, library usage, and operation sequences rather than simple pattern matching. We defined ten action categories: Data Exploration (examining distributions and patterns), Study Design Analysis (evaluating experimental conditions), Multi-source Integration (combining experimental and computational data), Statistical Analysis (hypothesis testing), Effect Size Calculation (biological significance assessment), Debugging (data validation), Group Statistics Calculation (summary computations), Results Export, Visualization, and Other Operations. To characterize workflow dynamics, we extracted action sequences from each conversation, compressed consecutive identical actions, and computed transition probabilities between action categories (n=11,659 sequences after filtering sequences <5 steps). We constructed directed graphs representing action transitions and analyzed temporal positioning of actions within workflows using relative position metrics (early: <25%, mid: 25-75%, late: >75%). Network analysis revealed transition patterns and workflow stages, with edge weights representing transition frequencies (minimum threshold: 100 transitions). All visualizations were generated using NetworkX (v2.8) and Matplotlib (v3.7), with statistical analyses performed in Python 3.9.

The RAG architecture provides comprehensive lifecycle management for document collections. This framework leverages asynchronous parallel processing for enhanced throughput, hybrid retrieval methods combining semantic and keyword searches for improved relevance, and hierarchical recall techniques to efficiently surface information from diverse granularities. Information retrieval is powered by these robust search capabilities, ensuring precise contextual data acquisition before generation. This design delivers a powerful and scalable RAG solution that effectively balances high performance with economic viability, ensuring flexible integration into diverse applications.

The RAG code was implemented in Python using custom scripts. The source code is publicly available on GitHub (https://github.com/Ard-Skelling/cheap-RAG, version 0.1.0) and detailed documentation is provided in the repository’s README file.

### Literature collection

To construct the specialized corpus for this study, we developed a systematic data ingestion pipeline to programmatically collect scholarly literature. The discovery phase targeted articles related to “aging” and “longevity” using the Semantic Scholar API, with a publication date range set between January 2020 and June 2025. This initial query identified a cohort of 20,872 relevant publications.

Our pipeline was designed for modularity and resilience. For each article, we extracted key metadata: title, authors, DOI, and publication venue. These entities and their intrinsic relationships were modeled and persisted as a knowledge graph within a Neo4j database. This graph structure captures the complex network of authors, papers, and journals in the target domain.

The acquisition of full-text articles was managed by a Redis-based task queue, which orchestrated the retrieval workflow. The system prioritized fetching documents from publicly accessible, open-access sources such as PubMed Central and BioMed Central. A multi-step retrieval process was employed.

All successfully retrieved full-text articles were archived in a MinIO object storage system for persistent storage. From the initial set of identified papers, this systematic process yielded a final corpus of 6,424 full-text articles. This curated collection formed the foundational dataset that was subsequently vectorized and ingested into our Retrieval-Augmented Generation (RAG) knowledge base for analysis.

### Intervention scoring methodology

The composite scoring system evaluated interventions using a systematic framework designed to identify promising therapeutic candidates while minimizing false positives from technical artifacts or biologically implausible findings. The scoring algorithm incorporated a critical plausibility filter followed by five weighted evaluation dimensions, producing scores ranging from 0 to 100 points with potential bonuses or penalties based on biological plausibility. To keep this score orthogonal to the statistical analysis result, we deliberately excluded p-values from the scoring criteria.

The plausibility assessment served as a primary quality control mechanism, applying severe penalties (-50 points) to interventions with fundamental biological implausibility, including age-related diseases erroneously presented as anti-aging interventions, extreme effect sizes without plausible mechanisms (beyond 3 times the interquartile range from quartile boundaries), results contradicting well-established aging biology, or biologically meaningless comparisons. Conversely, interventions demonstrating exceptional biological plausibility through targeting of established aging pathways with strong mechanistic rationale received bonus points (+30 points).

Experimental model quality assessment (0-25 points) stratified studies based on biological relevance and translational potential. Immortalized or cancer cell lines received no points, reflecting their limited relevance to physiological aging processes. Primary cell cultures earned 15 points, organoids and 3D culture systems 18 points, ex vivo tissue models 20 points, and *in vivo* studies using mammalian models or humans received the maximum 25 points. This hierarchical scoring prioritized interventions validated in physiologically relevant systems.

Aging biology relevance evaluation comprised two components: pathway involvement (0-15 points) and disease association (0-10 points). Pathway scoring assessed direct or indirect modulation of established aging mechanisms based on literature (e.g., mTOR signaling, sirtuins, NAD+ metabolism, autophagy, cellular senescence, and telomere maintenance)^2^. Disease association scoring evaluated connections to established age-related pathologies (e.g., Alzheimer disease, Parkinson disease, cardiovascular disease, type 2 diabetes, cancer)^40^.

Experimental rigor assessment (0-10 points) evaluated study design quality, considering the presence of appropriate controls, randomization procedures, blinding protocols, adequate sample sizes, and robust statistical analyses. Studies with poor design exhibiting major confounding factors received minimal scores, while those demonstrating rigorous experimental design with proper controls earned maximum points.

Translational potential evaluation (0-5 points) assessed human relevance and clinical feasibility. Interventions lacking human safety data received no points, those with limited human data or significant safety concerns earned 2 points, while interventions with established human use profiles and acceptable safety received maximum scores. This criterion prioritized candidates with realistic paths to clinical translation.

Novelty assessment comprised literature saturation (0-15 points) and mechanistic innovation (0-20 points) components. Literature saturation scoring inversely correlated with publication volume in aging research: well-established interventions with 50+ aging studies received no points, moderately studied interventions (10-50 publications) earned 3 points, emerging interventions (3-10 studies) received 10 points, and novel or repurposed interventions with fewer than 3 aging-specific studies earned maximum points. Mechanistic novelty scoring evaluated the discovery of novel aging mechanisms or unexpected findings, with paradigm-shifting discoveries receiving maximum scores.

### GenAge and DrugAge consistency analysis

To assess concordance between agent-derived intervention effects and established longevity annotations, we performed systematic benchmarking against the GenAge and DrugAge databases. For genetic perturbations, we evaluated the complete set of 796 significant and high-confidence interventions that met both filtering criteria: a false discovery rate below 0.05 and a final evaluation score of at least 80. Gene names were matched to GenAge entries using case-insensitive string searches across curated perturbation descriptors, including knockout, knockdown, deletion, deficiency, mutation, null, knockin, overexpression, rescue, and transgenic constructs. All matched entries were deduplicated at the gene level. For each matched intervention, we compared the inferred effect on biological age with the GenAge annotation of pro-longevity or anti-longevity. An intervention was considered concordant when its inferred direction of effect on biological age matched the expected direction based on the GenAge classification. Statistical significance of overall agreement was assessed with a one-sided binomial test using a null probability of 0.5. All matched cases were manually reviewed to confirm correct mapping between the experimental perturbation described in the original study and the corresponding GenAge annotation.

For pharmacological interventions, we applied the same filtering thresholds and evaluated all 279 significant and high-confidence compounds against DrugAge using exact and case-insensitive name matching. Directional agreement between agent-inferred effects and DrugAge annotations was assessed using the same criteria as in the genetic analysis. The one-sided binomial test with a null probability of 0.5 was used to determine significance of agreement.

### Genetic intervention type classification

To determine whether specific genetic intervention strategies differ in their effectiveness for anti-aging, we classified all significant genetic interventions (FDR < 0.05) by type using keyword matching. Interventions were categorized as: (1) Knockout: keywords including “knockout”, “knock-out”, “ko”, “null”, “deletion”, “deleted”, “-/-”; (2) Knockdown: keywords including “knockdown”, “knock-down”, “kd”, “shrna”, “sirna”, “rnai”; (3) Overexpression: keywords including “overexpression”, “overexpress”, “transgenic”, “tg”, “oe”; (4) Mutation: keywords including “mutant”, “mutation”, “mut”, “variant”.

For each intervention type, we calculated the proportion of interventions showing anti-aging effects (effect size < 0). We calculated the overall baseline anti-aging proportion from all significant genetic interventions and tested each intervention type against this baseline using binomial tests. Pairwise comparisons between intervention types were performed using Fisher’s exact test to compare the proportions of anti-aging versus pro-aging interventions. P-values were adjusted for multiple testing using the Benjamini-Hochberg false discovery rate correction. Effect size distributions for each intervention type were visualized using kernel density estimation.

### Pathway enrichment analysis

To validate the biological plausibility of identified age-modifying genetic interventions, we performed Over-Representation Analysis (ORA) using the GSEApy Enrichr API (v1.0+). Gene names were extracted from intervention condition names using keyword matching and validated against official gene reference databases. We removed common suffixes (knockout, overexpression, etc.) and matched extracted names to mouse gene symbols from KEGG_2019_Mouse and GO_Biological_Process_2021 pathway databases, supplemented with MyGene.info queries.

ORA was performed using the validated significant genes (FDR < 0.05) as the query set and all genes from genetic interventions as the custom background. We tested for enrichment in KEGG_2019_Mouse pathways using one-sided Fisher’s exact test with automatic Benjamini-Hochberg FDR correction. Pathways were filtered to include only those with pathway size < 100 genes and overlap ≥ 15 genes to focus on well-characterized, substantially enriched pathways. Enrichment significance was set at FDR < 0.05. Cancer-related pathways are filtered out for visualization purposes.

### FDA-approved drug clinical validation

To assess clinical relevance and translational potential, we classified significant drug interventions (FDR < 0.05) by clinical development stage using the OpenTargets drug database^21^. Drug names were matched to OpenTargets entries through multiple strategies: (1) exact name matching, (2) synonym and trade name matching, (3) token-based matching for combination drugs, (4) fuzzy string matching with validation rules (similarity threshold ≥ 0.80, prefix matching, exclusion of descriptor mismatches), and (5) conservative substring matching.

Drugs were classified into clinical development stages based on OpenTargets annotations: FDA Approved (isApproved = True or maximumClinicalTrialPhase = 4.0), Phase 3 (maximumClinicalTrialPhase = 3.0), Phase 2 (maximumClinicalTrialPhase = 2.0), Phase 1 (maximumClinicalTrialPhase = 1.0), and Preclinical (Phase < 1 or no phase information). For each clinical stage, we calculated the proportion of anti-aging interventions (log2FC < 0) and compared proportions across stages using chi-square test. Pairwise comparisons between stages were performed using Mann-Whitney U tests with FDR correction. We additionally tested each stage against the overall baseline using Fisher’s exact test. Top FDA-approved anti-aging drugs were ranked by effect size and annotated with mechanisms of action based on drug type and pharmacological class information from OpenTargets.

### Quality control and validation

Multiple quality control steps ensured analytical accuracy. Datasets with fewer than 3 samples per group were excluded. For the final analysis, we applied stringent outlier filtering using the interquartile range (IQR) method with a 10× threshold. Specifically, we calculated Q1 and Q3 for both beta coefficients and -log10(p-values), then removed data points outside [Q1 - 10×IQR, Q3 + 10×IQR]. This filtering removed 619 outliers (1.4% of 44,148 initial comparisons) that likely represented technical artifacts, data errors, or extreme biological states incompatible with standard aging trajectories.

Agent decisions underwent rigorous validation through manual review of 100 randomly selected analyses by two independent doctoral-level bioinformaticians with expertise in aging research and statistical genomics. Reviewers assessed each analysis across eight standardized criteria: (1) hypothesis formulation quality, requiring clear identification of control versus treatment groups; (2) appropriateness of statistical test selection for the experimental design and data structure; (3) accuracy and completeness of p-value calculations; (4) correctness and reasonableness of effect size computations; (5) adherence to protocol restrictions against direct chronological age comparisons; (6) analytical completeness, including presence of all required result columns; (7) successful integration of metadata and aging clock data; and (8) conformity to the specified analytical workflow.

Each criterion received a binary score (1 = correct, 0 = issue identified) along with descriptive notes for any deviations. To accurately assess agent performance versus data quality, reviewers distinguished between genuine agent errors (e.g., inappropriate statistical test, forbidden comparison) and data-driven limitations (e.g., insufficient sample size, extreme effect sizes, file parsing failures). Data limitations received scores of 0 with explanatory notes but were excluded from correctness rate calculations, as these issues originate from the input datasets rather than agent logic.

### Interactive platform development

The ClockBase web interface was developed using React and FastAPI, providing real-time access to all clock predictions and analytical results. The platform implements three analysis modules: group comparison for case-control studies, correlation analysis for continuous variables, and accuracy assessment for clock validation. Users can upload new datasets for instant biological age calculation through Biolearn’s integrated preprocessing pipelines, which handle quality control, probe filtering, normalization, and missing value imputation automatically. The backend leverages Biolearn’s Python API for clock pre-calculations, ensuring consistency with the validated implementations used in the original publications. All data and results are downloadable in standard formats to facilitate downstream analysis. The platform is hosted on cloud infrastructure with automatic scaling to handle concurrent user sessions, with Biolearn’s efficient computation enabling real-time calculation of multiple aging clocks simultaneously.

### Transcriptomic age analysis of ouabain effect

Raw counts from GSE122080^23^ were filtered to retain only genes with ≥10 reads in at least 20% of samples. Filtered RNA-seq data were then processed by applying Relative Log Expression (RLE) normalization, log transformation, and YuGene normalization^41^. Missing expression values for clock genes absent from the dataset were imputed using their precomputed average values. Normalized expression profiles were subsequently centered to the median profile of young control samples.

tAge for each sample was estimated using Bayesian Ridge-based multi-tissue tAges of chronological age and expected mortality, trained on a large collection of independent datasets that excluded GSE122080^14^. Pairwise differences in tAge between groups were evaluated with a mixed-effects model, incorporating both the mean and standard error of tAge predictions. P-values were adjusted for multiple comparisons using the Benjamini-Hochberg (BH) procedure^42^.

To estimate the contribution of individual genes to ouabain-induced tAge changes in old animals, we multiplied each gene’s log fold change (logFC) between old control and ouabain-treated mice by its corresponding clock coefficient. Statistical significance of gene contributions was derived from differential expression analysis performed with limma package in R^43^.

Module-specific tAges of chronological age and expected mortality, trained with Elastic Net models on independent datasets, were applied to the scaled relative expression profiles within the same framework. Resulting tAge estimates from each module-specific clock were standardized across all old samples (control and ouabain-treated). For each module clock, standardized tAge was compared between groups using ANOVA, and p-values were adjusted for multiple testing using the BH method.

### Animals

All experiments were conducted following NIH guidelines and protocols approved by the IACUC at Brigham and Women’s Hospital (BWH), animal protocol 2019N000042 (Characterization of mouse physiology, metabolism, and aging). All mice were housed in Allentown standard ventilated cages at 22–23.5 °C (room temperature) with 40–70% humidity on a 12-h light/dark cycle, with standard mouse chow diet (5053 PicoLab® Rodent Diet 20) and water provided ad libitum. Cages were maintained by facility staff and changed biweekly or earlier if soiled. 20-month-old male C57BL/6 mice were obtained from the NIA Aging Colony and randomized into control and experimental groups based on baseline frailty index scores within cages, minimizing potential bias due to differences in chronological age among non-littermates.

### Ouabain injections

Ouabain octahydrate, ≥95% (HPLC) powder, was purchased from Sigma-Aldrich, O3125-250MG, CAS: 11018-89-6. A stock solution of ouabain was prepared by dissolving 3,000-4,000 µg of ouabain in 27,000–36,000 µL of PBS (pH 7.4; Thermo Scientific, 10010023). Following weight-based dose calculation, mice were administered the appropriate volume of ouabain solution via intraperitoneal injection using 1 mL syringes (BD 309628) and sterile 26-gauge veterinary hypodermic needles (Air-Tite, N2612). Ouabain injections were administered twice per week, every third week, over a period of three months. Control mice received weight-adjusted amounts of PBS.

### Frailty Index

Frailty Indexes were assessed as described by Whitehead et al.^24^, with the exception that body temperature was not measured. Assessments were performed prior to treatment initiation (at 20 months of age) and after three months of treatment in both control and ouabain-injected mice. Differences in frailty index and individual frailty parameters between groups at a given age were assessed using the Wilcoxon rank-sum test. P-values for individual parameters were adjusted for multiple comparisons using the BH method. Longitudinal changes in frailty index over the treatment period within each group were evaluated with the Wilcoxon signed-rank test.

### Body composition

Body composition was assessed in conscious mice using the EchoMRI 3-in-1 system (EchoMRI, Houston, TX), acquired with support from NIH S10 grant S10OD032157. Mice were briefly acclimated (∼2 minutes) to the plastic tube without restraint, after which the restraint was applied and mice were positioned in the scanner for data acquisition. Group differences in body mass and fat-to-lean ratio at a given age were assessed using the Wilcoxon rank-sum test. Longitudinal changes in these measures within each group over the treatment period were evaluated using the Wilcoxon signed-rank test.

### Echocardiography

Echocardiographic measurements were performed using a FUJIFILM VisualSonics Vevo 3100 ultrasound system equipped with the MX550S high-frequency linear array transducer operating at 40 MHz with 100% power, 14 fps frame rate, and 31 dB gain. Mice were anesthetized and positioned supine on a heated imaging platform to maintain body temperature, with heart rate, respiratory rate, and body temperature continuously monitored throughout the procedure. The chest area was shaved, and ultrasound gel was applied to ensure optimal acoustic coupling. Two-dimensional and M-mode echocardiographic images were acquired from parasternal long-axis views using the Mouse (Small) Cardiology PSLAX preset application, with dynamic range set to 60 dB, C5 display map, gate depth of around 9 mm, and gate size of around 7 mm for optimal visualization of cardiac structures. Cardiac output, stroke volume, left ventricular internal dimensions in diastole (LVIDd) and systole (LVIDs) were calculated using the VisualSonics measurement package algorithms. All measurements were performed by a single experienced operator analyzing a minimum of six cardiac cycles for each parameter, with average values recorded and data exported from the VisualSonics system for further analysis, ensuring image quality assessment and clear endocardial border definition before inclusion in the analysis. Mean cardiac output and its standard error were calculated for each animal across echocardiography runs. Differences in cardiac output between 23-month-old saline- and ouabain-treated male mice were assessed using a mixed-effects model. The same statistical approach was applied to evaluate differences in stroke volume, LVIDs, and LVIDd. Resulting p-values were adjusted for multiple comparisons using the BH method.

### Brain sectioning and immunostaining

Brains were dissected in cold phosphate-buffered saline (PBS) and bisected longitudinally to separate the two hemispheres. The right hemispheres were fixed overnight at 4°C in 4% paraformaldehyde (PFA) with gentle rotation. Following fixation, tissues were cryoprotected in 30% sucrose overnight at 4°C, then embedded in OCT compound and frozen. Sagittal brain sections (35 µm thick) were obtained using a cryostat. Sections were blocked for 1 hour at room temperature in 5% bovine serum albumin (BSA) with 0.5% Triton X-100 in PBS. For microglia staining, sections were incubated overnight at 4°C in a humidified chamber with rabbit anti-Iba1 primary antibody (1:400, Wako 019-19741, FUJIFILM Irvine Scientific) diluted in PBS containing 0.25% Triton X-100. After primary antibody incubation, sections were washed three times for 10 minutes each in 1× PBS, followed by incubation with Alexa Fluor 647-conjugated anti-rabbit secondary antibody (1:500, Thermo Fisher Scientific, A21244, Lot 2659317) and DAPI (1:1000, Sigma-Aldrich, D9542) diluted in PBS with 0.1% Triton X-100. Slides were mounted with ProLong™ Glass Antifade Mountant (Invitrogen, P36980, Lot 2801574) and stored at 4°C overnight prior to imaging.

### Imaging and microglia morphology quantification

For each brain, three images of the hippocampal CA1 region and three images of the dorsal cortex were randomly selected and acquired using a Nikon Ti2 inverted spinning disk confocal microscope equipped with an Andor Zyla 4.2 Plus sCMOS monochrome camera. All images were obtained using a Plan Fluor 40×/1.3 NA oil immersion DIC H/N2 objective. Imaging parameters, including laser intensity, exposure time, and threshold, were kept constant across all samples to ensure comparability. Images were processed in ImageJ to generate 2D projections and to apply linear adjustments to brightness and contrast. Microglial morphology was quantified by segmenting the images using the ImageJ plugin MicrogliaMorphology^44^. For every region and mouse, we calculated mean number of branches (in log-scale), mean number of junctions (in log-scale), mean maximum branch length (in log scale), and mean total area occupied by microglia cells (in log scale) across sections, together with the corresponding standard errors. Differences in these parameters between control and ouabain-treated groups were assessed using mixed-effects models separately for each region. Resulting p-values were adjusted for multiple comparisons using the BH method.

### Indirect calorimetry

Metabolic rate and activity were assessed at the Brigham and Women’s Hospital Metabolic Core Facility. Oxygen consumption (VO2), carbon dioxide production (VCO2), respiratory exchange ratio (RER), food intake, and ambulatory and locomotor activity were measured using a Comprehensive Lab Animal Monitoring System (CLAMS; Columbus Instruments, Columbus, OH). Mice were individually housed in sealed metabolic cages and acclimated for 14–20 hours prior to data collection to minimize stress-related artifacts. Data were recorded over a 48-hour period under standardized conditions (12-hour light/dark cycle; ambient temperature 22.5 °C) with ad libitum access to food and water. Gas sensors were calibrated according to the manufacturer’s instructions. Negative food intake values were excluded from the resulting dataset, and hourly averages for all measured metabolic parameters, as well as calculated energy expenditure and energy balance values, were obtained using CalR, a web-based tool for indirect calorimetry analysis^45^. An additional group of middle-aged C57BL/6J mice (400 days old) was included to establish reference values. Indirect calorimetry data were analyzed using statistical models that accounted for repeated measures and body weight (for weight-dependent variables). Group-level differences for weight-dependent physiological variables were assessed by ANCOVA with body weight as a covariate, while weight-independent outcomes were analyzed by one-way ANOVA. Pairwise comparisons were performed between old saline-treated mice and both middle-aged reference mice and old ouabain-treated mice, with p-values adjusted using the BH method. To account for repeated measurements, a mixed-effects model was applied with body weight and time as fixed effects and mouse ID as a random effect. Analyses were conducted separately for dark and light phases, and only significant or marginally significant pairwise effects (adjusted p-value < 0.1) are shown in the figures.

## Discussion

ClockBase represents a paradigm shift in aging research methodology, demonstrating how specialized AI agents can systematically mine previous molecular datasets to accelerate therapeutic discovery. This approach transforms decades of archived samples, originally collected for unrelated purposes, into a discovery engine for age-modifying interventions. The identification of thousands of significant age-modifying effects, including unexpected candidates like ouabain alongside established longevity drugs, validates the power of hypothesis-free discovery at scale.

Our results demonstrate the practical utility of ClockBase for aging research. The platform successfully integrates population-scale molecular data with AI-guided analysis to systematically identify age-modifying interventions. To validate this approach, we experimentally tested ouabain, a top-scoring candidate not previously studied for its anti-aging potential. The compound produced broad geroprotective effects in aged mice, including reduced frailty progression, improved cardiac function, and decreased neuroinflammation. This proof-of-principle validation confirms that ClockBase can prioritize compounds with genuine therapeutic potential from large datasets. Translation of any candidate to humans will require careful safety evaluation, robust multi-tissue biomarker endpoints, and thorough mechanistic understanding. The concordance between clock predictions and functional outcomes supports the utility of biological clocks as discovery tools while emphasizing the necessity of rigorous experimental validation.

The success of the AI agent system in achieving analytical accuracy across diverse experimental contexts demonstrates that current language models, when properly specialized and validated, can perform complex bioinformatic analyses with expert-level competence. The agent’s ability to select appropriate statistical methods, handle edge cases, and generate biologically meaningful interpretations suggests that AI-assisted research can dramatically accelerate the pace of discovery while maintaining scientific rigor. Notably, the multi-dimensional scoring system successfully prioritized interventions that balance biological plausibility, novelty, and translational potential, addressing a key challenge in drug discovery.

The bimodal distribution of intervention effects, with distinct distributions for age-accelerating and age-reversing perturbations, suggests that the AI agent successfully detects signals in both directions. Among filtered data, genetic interventions comprised the largest category (46.0%) though with modest significance rates (10.0%), underscoring the polygenic complexity of aging regulation. Surprisingly, disease states showed the highest significance rate (24.3%), followed by environmental exposures (15.2%) compared to drug treatments (11.3%), highlighting both the impact of pathology on aging and the untapped potential for environmental and lifestyle interventions. After stringent outlier filtering, effect sizes remained substantial (ranging from - 23.12 to 24.35) while being more biologically plausible than the extreme values initially observed.

The platform addresses several critical limitations in current aging research. First, the systematic reanalysis through an aging lens reveals insights missed by original studies focused on other phenotypes. Second, for DNA methylation data, Biolearn’s harmonized processing pipeline enables direct comparison across studies, while for RNA-seq data, the standardized preprocessing ensures consistent application across datasets. Third, the AI agent’s consistent analytical framework eliminates subjective interpretation bias. Fourth, the comprehensive scoring system moves beyond p-values to identify truly promising interventions. The use of diverse clock types, including Biolearn’s methylation clocks for epigenetic aging, the mortality predictor for mouse transcriptomic data, and the multi-species clock for human RNA-seq, provides multiple complementary perspectives on biological aging. However, important caveats remain: different clock types capture distinct aspects of aging (chronological vs. biological vs. mortality risk vs. rate of change); mouse findings require careful translation to humans; and the reliance on public data introduces unequal data quality and selection bias toward well-studied interventions, while underrepresenting novel or less-characterized ones.

Critically, multiple systems-level analyses validate that the identified effects represent genuine biological patterns rather than technical artifacts. The observation that disease perturbations show both the highest significance rate (24.3%) and strong directional bias toward age acceleration (83% pro-aging) provides internal validation, as pathological states are universally expected to accelerate aging. The systematic hierarchy within genetic intervention types, where loss-of-function approaches consistently outperform gain-of-function approaches (1.85-fold enrichment for knockdown, 1.45-fold for knockout vs overexpression), reflects established biological principles of cellular homeostasis and argues against random technical variation. The convergence of age-modifying genes on canonical aging pathways (cellular senescence, longevity regulation, autophagy) through independent pathway enrichment analysis demonstrates that diverse interventions tested across thousands of studies point to the same fundamental biology. Finally, the significant concordance with curated longevity databases (81% for GenAge, 100% for DrugAge), compiled from completely independent lifespan studies using different organisms and methods, provides the strongest cross-platform validation. These multi-level, orthogonal validations collectively demonstrate that ClockBase Agent captures coherent biological aging signals that are robust to technical artifacts, batch effects, and study-specific biases.

The overall positive skew in effect sizes, where interventions more commonly accelerate than decelerate aging, reveals a fundamental biological asymmetry. This pattern appears consistently across all intervention categories and reflects a core principle: disrupting optimized biological systems is easier than improving them. Disease states produce the largest and most consistent aging effects, environmental perturbations show intermediate impacts, while genetic interventions display the highest redundancy and compensation. This hierarchical sensitivity to perturbation types would not emerge from technical artifacts and validates that aging clocks measure genuine biological vulnerability rather than experimental noise. The finding that most FDA-approved drugs show pro-aging effects (136 vs 78 anti-aging, p=0.0001) further suggests that current therapeutics, while treating specific conditions, may inadvertently accelerate biological aging as a side effect.

Our experimental validation of ouabain, a top-scoring AI-identified candidate, demonstrates the practical utility of ClockBase for discovering genuine geroprotectors. Intermittent ouabain treatment in aged mice produced broad improvements across multiple aging phenotypes, including halted frailty progression, enhanced cardiac output, and reduced neuroinflammation. The frailty index, which integrates multiple phenotypes and correlates with health and mortality in both mice and humans^24,46,47^, showed no increase over three months in ouabain-treated animals compared to significant progression in controls, with improvements across multiple domains without adverse effects. These proof-of-principle results confirm that ClockBase can prioritize compounds with genuine therapeutic potential, demonstrating that transcriptomic age predictions translate into tangible health benefits.

Looking forward, ClockBase provides a foundation for precision longevity medicine. The platform enables researchers to query how specific interventions affect biological age across diverse genetic backgrounds, ages, and tissue types. Integration with single-cell and spatial sequencing data will reveal cell-type-specific aging trajectories. Incorporation of human cohort studies will validate findings across species. Most importantly, the open-access design democratizes aging research, allowing any investigator to test hypotheses against millions of samples instantly using validated aging biomarkers.

The broader implications extend beyond aging research. The successful deployment of specialized AI agents for systematic knowledge extraction from public databases could transform how we approach biological discovery. Rather than the traditional hypothesis-driven model where researchers analyze individual datasets, AI agents can continuously scan the entirety of published data, identifying patterns invisible to human analysis. This approach is particularly powerful for complex, multifactorial processes like aging where subtle signals may emerge only from massive sample sizes. As biological databases grow exponentially, AI-mediated discovery will become essential for extracting actionable insights from the data deluge.

In conclusion, ClockBase demonstrates that combining comprehensive data integration, standardized aging biomarkers, and specialized AI analysis can accelerate the identification of longevity interventions. By making biological age measurable across millions of samples through diverse biomarkers, we enable a new era of data-driven longevity research. The platform’s discoveries, from validating known longevity drugs to identifying novel candidates like ouabain, showcase the potential for AI-human collaboration to solve one of biology’s most fundamental challenges. As we continue expanding with new data types and analytical capabilities, ClockBase will evolve into essential infrastructure for the emerging field of longevity medicine.

## Acknowledgments

We thank members of Biomarkers of Aging Consortium for their valuable feedback and suggestions. We thank the members of the Biolearn team for providing tools for aging biomarker harmonization. We thank OpenAI and Avinasi Labs for supporting ChatGPT and Claude API costs. K.Y. is supported by the NIH/NIA F99/K00 award (F99AG088431 and K00AG088431). Supported by grants to V.N.G. from NIA and Hevolution.

## Author contributions

K.Y. conceptualized and initiated the study, collected the data. K.Y and A.T. performed data analyses. A.T. applied RNA clock and RNA preprocessing. H.W. performed agent analysis. A.M., C.G.D.M., S.I., A.E.G., A.T., and J.R.P. conducted *in vivo* mouse model validation of ouabain. K.Y., C.L., K.H., and Y.Q. designed the AI agent system. K.Y., A.T., and D.G. designed the website. C.J. developed the literature RAG system. D.L. and H.L. performed agent trace evaluation. L.P.D.L.C. and C.K. contributed to DNA methylation clock prediction. A.Z. supported agent analysis and writing. K.Y., J.M.V.R., L.C., A.R., J.L., T.W.-C., and V.N.G. supervised the study and contributed to paper writing. V.N.G. obtained funding and provided the overall leadership. All authors contributed to the design of the study and writing of the manuscript.

---

**Extended Data Figure 1.**
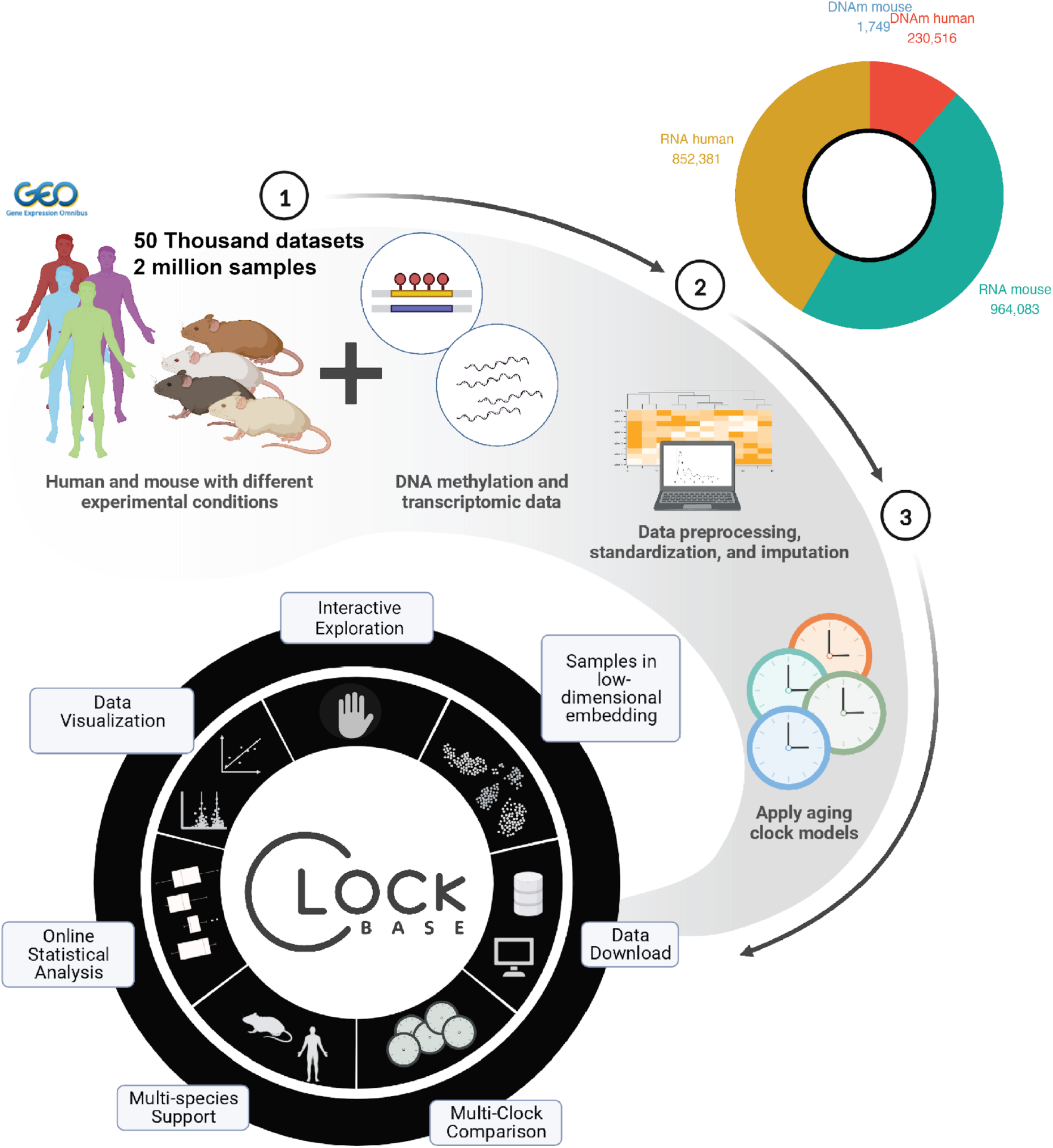
Schematic plot showing the design and data flow of ClockBase website, showing the data source and all key functionality of the website.

**Extended Data Figure 2.**
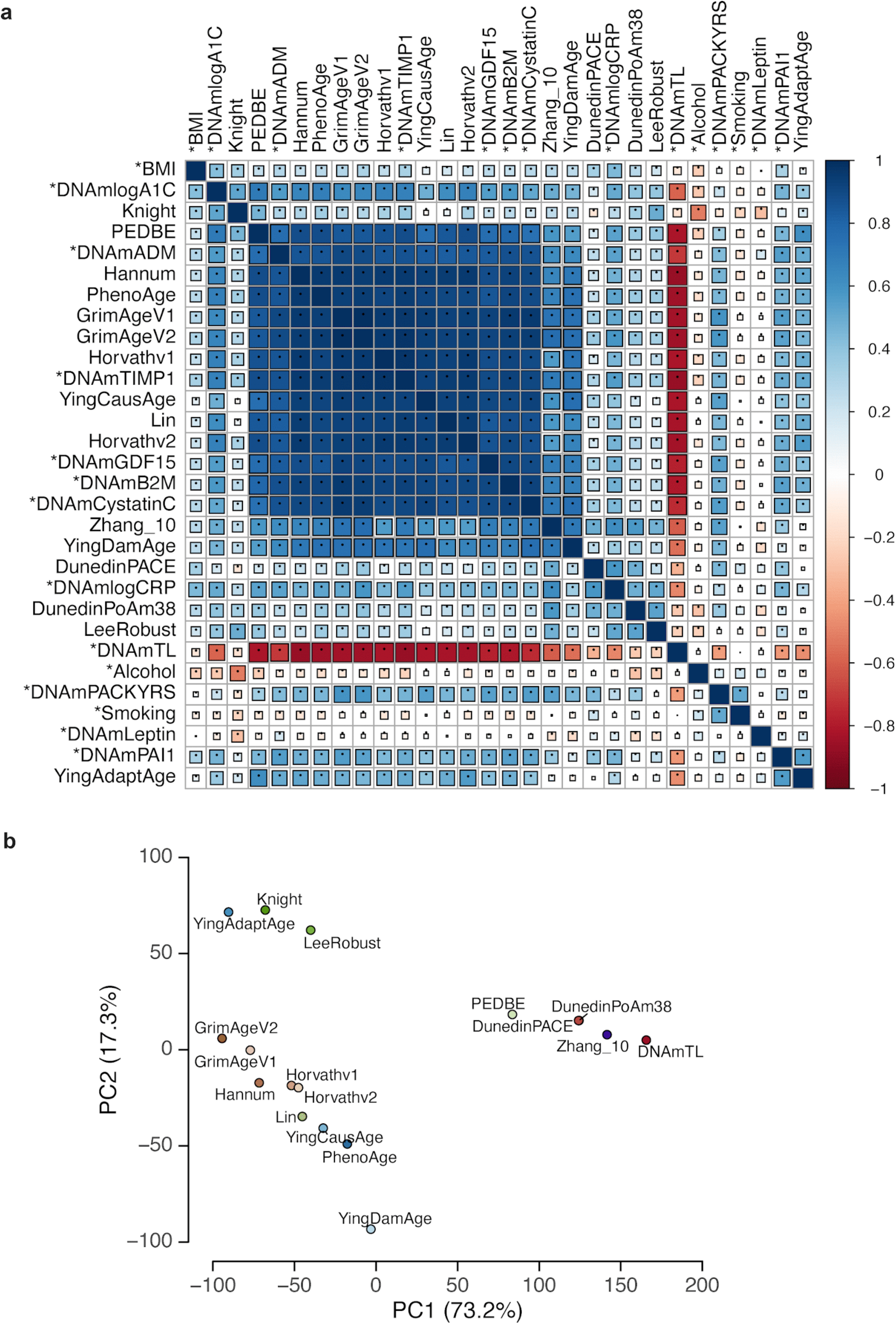
DNA methylation aging clock analysis in human blood samples. **a.** Correlation matrix heatmap displaying pairwise Pearson correlation coefficients between 39 epigenetic aging clocks calculated from human blood DNA methylation samples. Each cell represents the correlation between two clocks, with rows and columns ordered by hierarchical clustering. The color scale ranges from dark blue (correlation = -1, perfect negative correlation) through white (correlation = 0, no correlation) to dark red (correlation = 1, perfect positive correlation). Square size is proportional to the absolute correlation value, with larger squares indicating stronger correlations. Asterisks (*) denote clocks with established biological or clinical associations. **b.** Principal component analysis (PCA) plot showing the distribution of 17 blood epigenetic aging clocks based on their predictions across human blood DNA methylation samples. The x-axis represents the first principal component (PC1) and the y-axis represents the second principal component (PC2). Each point represents one aging clock.

**Extended Data Figure 3.**
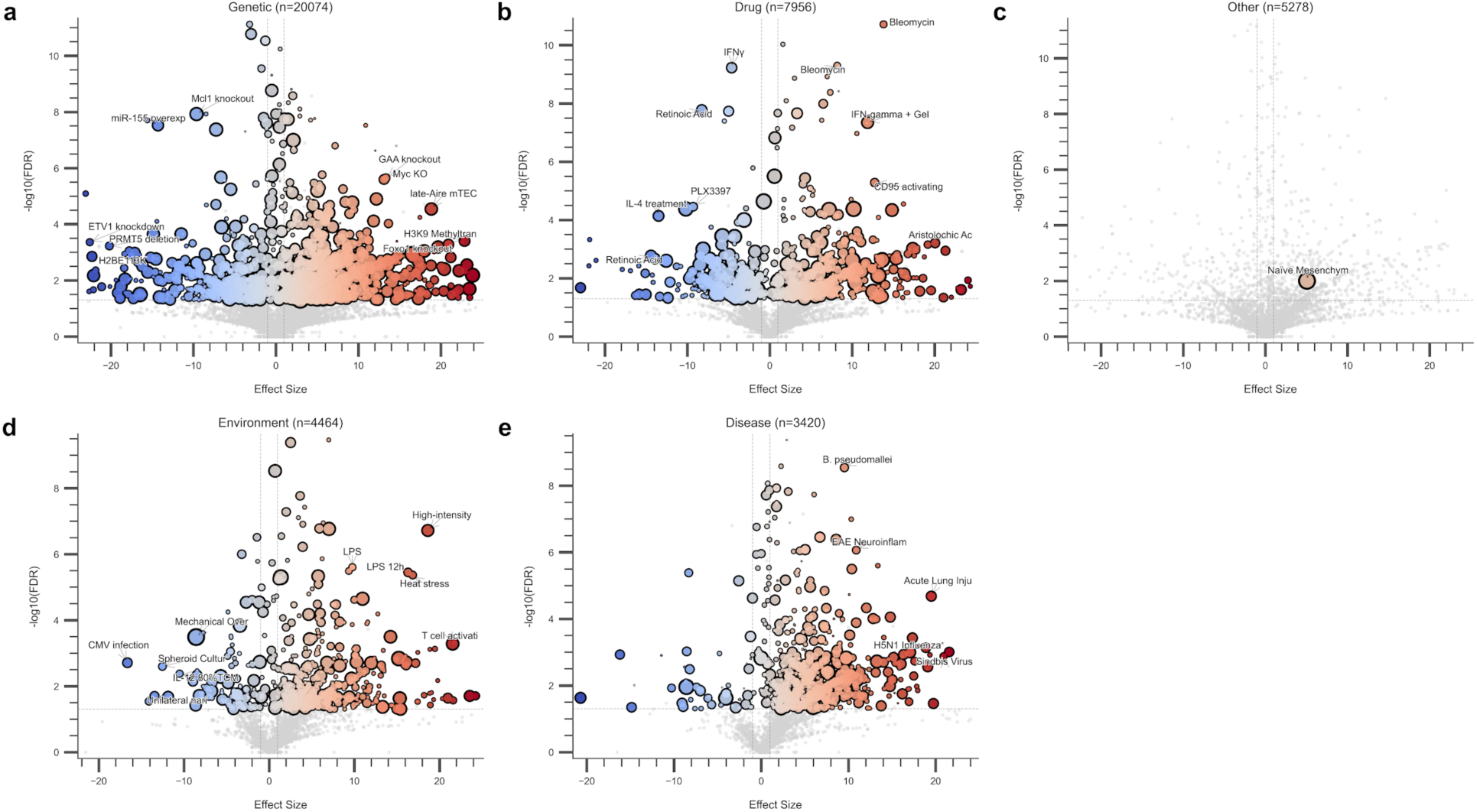
Volcano plots of age-modifying interventions across intervention categories. Each panel displays interventions analyzed by the AI agent framework, categorized as **a**, Genetic (n=20,074), **b**, Drug (n=7,956), **c**, Other (n=5,278), **d**, Environment (n=4,464), and **e**, Disease (n=3,420). The x-axis represents the effect size (predicted change in transcriptomic age), while the y-axis shows statistical significance as -log10(FDR). Blue circles indicate anti-aging interventions (negative effect size), red circles indicate pro-aging interventions (positive effect size), and gray circles represent non-significant interventions (FDR > 0.05). Circle size is proportional to the Final Score, with larger circles indicating more reliable results scored by the AI agent framework. Selected interventions with the strongest effects or highest significance are labeled. Horizontal dashed lines indicate FDR significance thresholds.

**Extended Data Figure 4.**
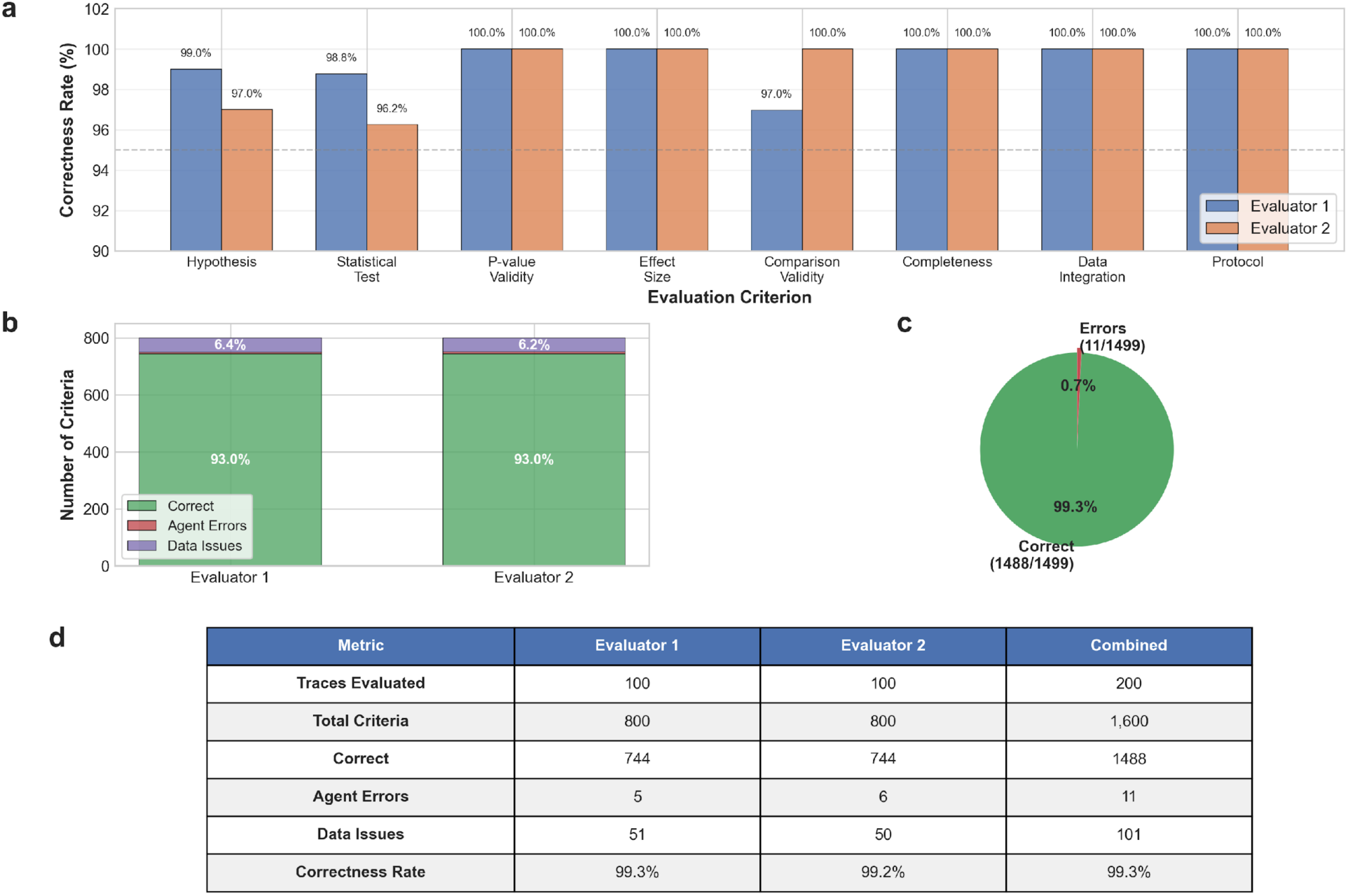
Manual validation of agent analytical accuracy. **a.** Correctness rates across eight evaluation criteria for two independent evaluators. Bar heights represent the percentage of correct assessments for each criterion: hypothesis formulation, statistical test selection, p-value validity, effect size calculation, comparison validity, analytical completeness, data integration, and protocol adherence. **b.** Distribution of assessment outcomes for each evaluator. Stacked bars show the proportion of correct assessments (green), agent errors (red), and data-related issues (purple) out of 800 total criteria evaluated per evaluator. **c.** Overall correctness rate across all assessments. **d.** Summary table of validation metrics for both evaluators and combined results, including number of traces evaluated, total criteria assessed, correct assessments, agent errors, data issues, and final correctness rates.

**Extended Data Figure 5.**
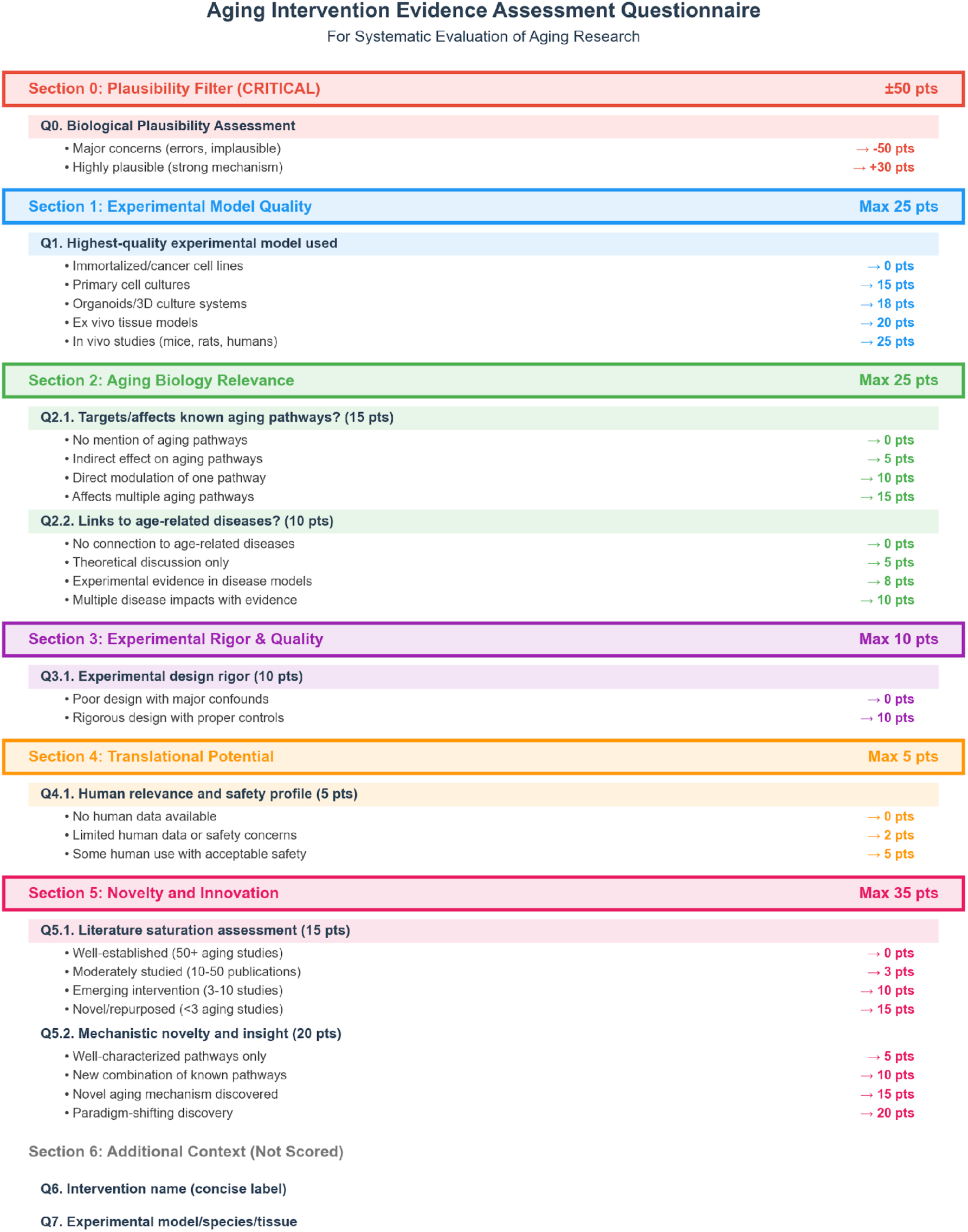
Multi-dimensional scoring system for prioritizing aging interventions. Systematic evaluation framework used by the AI agent to assess and rank interventions across eight criteria: biological plausibility filter (±50 points), experimental model quality (0-25 points), aging biology relevance (0-25 points), experimental rigor and quality (0-10 points), translational potential (0-5 points), and novelty/innovation (0-35 points). The system generates composite scores ranging from -50 to 140 points to prioritize candidates for experimental validation.

**Extended Data Figure 6.**
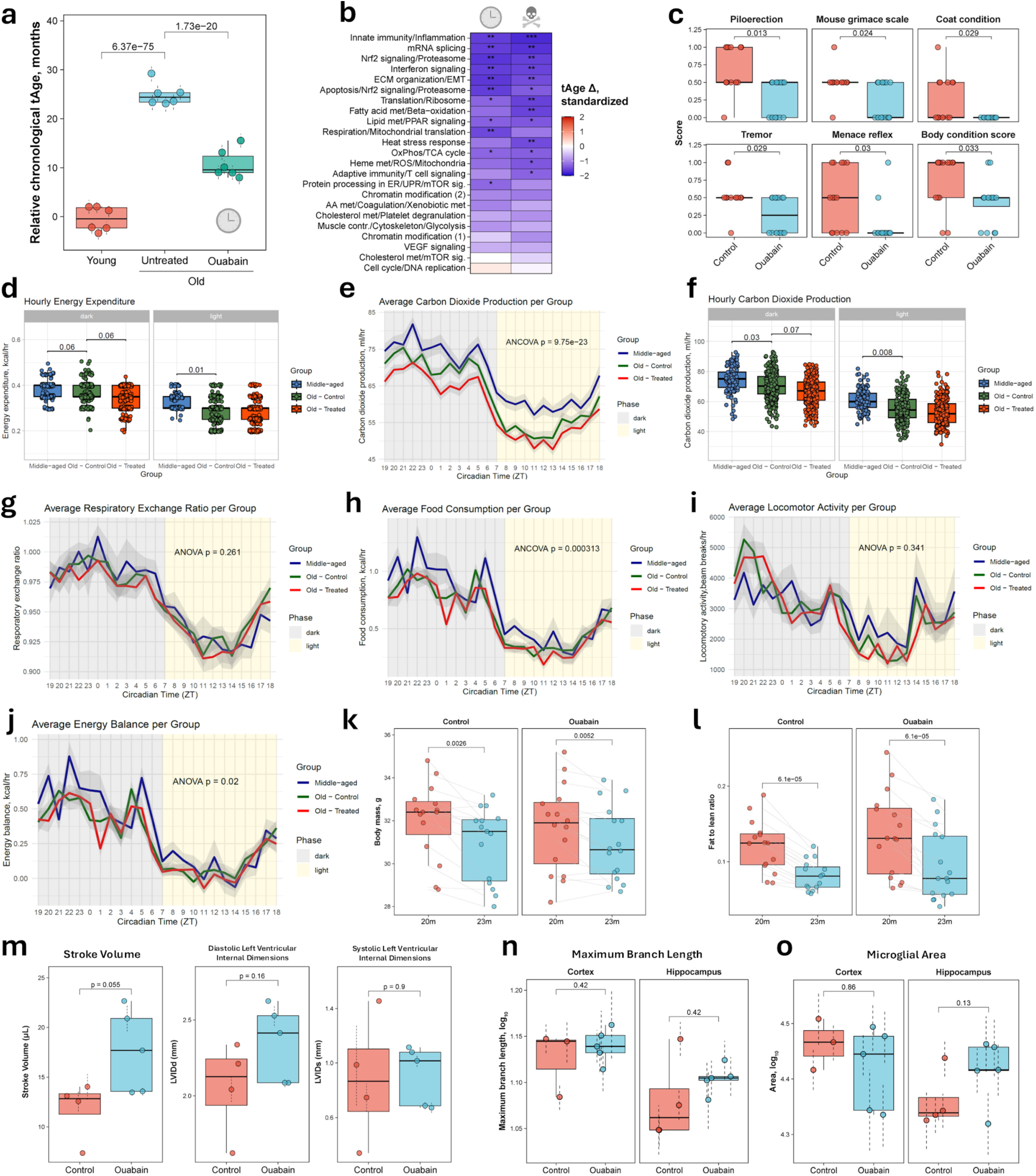
Ouabain’s effect on molecular, physiological, and clinical profiles of aged mice. **a.** tAge predicted by chronological rodent multi-tissue clock for young (7-week-old), old (24-month-old) untreated and ouabain-treated C57BL/6 female mice (n=6 per group). Data are mean tAges ± s.d. **b.** Standardized tAge differences induced by ouabain in old female mice according to module-specific rodent multi-tissue clocks of chronological age (left) and expected mortality (right). P-values were assessed with one-way ANOVA and adjusted for multiple comparisons with the BH method. * p.adj < 0.05; ** p.adj < 0.01; *** p.adj < 0.001. **c.** Comparison of representative frailty parameters between old control and ouabain-treated male mice after 3 months of injections. BH-adjusted p-values derived with the Wilcoxon rank-sum test are indicated in text. **d.** Comparison of hourly circadian energy expenditure between middle-aged and old control and ouabain-treated mice during dark (left) and light (right) phases. BH-adjusted p-values are indicated with text. **e.** Average carbon dioxide production over 24 hours (averaged 48-hour data) in middle-aged and old control and ouabain-treated male mice. The x-axis shows time in hours. Light and dark phases are depicted with the background color. Experimental groups are shown with line colors. Shaded bands around lines indicate standard error. **f.** Comparison of hourly carbon dioxide production between middle-aged and old control and ouabain-treated mice during dark (left) and light (right) phases. BH-adjusted p-values are indicated with text. **g-j.** Average respiratory exchange ratio (RER) **(g)**, food consumption **(h)**, locomotor activity **(i)**, and energy balance **(j)** over 24 hours (averaged 48-hour data) in middle-aged and old control and ouabain-treated male mice. The x-axis shows time in hours. Light and dark phases are depicted with the background color. Experimental groups are shown with line colors. Shaded bands around lines indicate standard error. **k-l.** Dynamics of body mass **(k)** and fat-to-lean ratio **(l)** over 3 months in old control (left) and ouabain-treated (right) male mice. Change over time was assessed for each group with the Wilcoxon signed-rank test. Error bars show standard error of the mean. Shaded areas indicate 95% confidence intervals. **m.** Comparison of stroke volume (left), diastolic left ventricular internal dimensions (DLVID) (center), and systolic left ventricular internal dimensions (SLVID) (right) assessed with echocardiography in old control and ouabain-treated male mice after 3 months of injections (n=4-5 mice per group). P-values derived with a mixed-effect model are shown in text. Data are mean ± s.e.m. **n-o.** Maximum branch length **(n)** and microglial area **(o)** in cortex (left) and hippocampus (right) of old control and ouabain-treated male mice after 3 months of injections (n=4-5 mice per group). P-values were derived with the mixed-effect model and adjusted for multiple comparisons with the BH method. Data are mean ± s.e.m. Boxplots: center line indicates median; boxes, interquartile range; whiskers, ±1.5× IQR.

